# Reconstructing the deep phylogeny of the MAPK signaling network: functional specialization via multi-tier coevolutionary expansion

**DOI:** 10.1101/2024.10.01.616093

**Authors:** EJ Huang, Jeeun Parksong, Amy F. Peterson, Fernando Torres, Sergi Regot, Gabriel S. Bever

## Abstract

The mitogen-activated protein kinase (MAPK) signaling network is a three-tier cascade that regulates key cellular responses in eukaryotes. However, the evolutionary origins of its complex interactions and functional diversity remain poorly understood. Here, we conducted a comprehensive phylogenetic analysis of MAPK components across Eukarya to delineate divergences of non-human orthologs of human paralogs along the human evolutionary backbone. We identified two major pulses of coevolutionary expansion: one predating the divergence of fungi and animals, and another predating the origin of animals. Our reconstruction also infers a polyphyletic origin for the atypical MAPKs. Integrating functional literature across eukaryotic taxa with our reconstructed trees reveals that the two clades of MAP3K, Sterile-like (STE) and tyrosine kinase-like (TKL), had distinct evolutionary trajectories and influences on downstream pathway diversification. STEs that function as MAP3Ks are conserved across extant eukaryotes. Despite the absence of TKL MAP3Ks in many early diverging eukaryotes, the expansion of TKL MAP3Ks aligns phylogenetically and functionally with that of the downstream MAP2Ks and MAPKs. We thus propose that the MAPK network originated as a STE-regulated pathway, and that subsequent radiations of the TKLs drove the diversification of downstream components and top-down finetuning of pathway specificity. We thus provide an evolutionary framework for generating novel hypotheses on the functional diversity of this key signaling network, including potential insights into the evolution of animal multicellularity. Our study demonstrates that phylogenetics can offer new perspectives towards understanding complex cellular physiology.

**Significance:** The mitogen-activated protein kinase (MAPK) signaling network responds to various signals and regulates basic cell physiology including proliferation and apoptosis. This three-tier network is universal in eukaryotes, but there is great functional diversity among different homologs as well as different taxa. Here we compared amino acid sequences to reconstruct MAPK evolutionary history as a network. We found that the three levels expand in parallel, showcasing a special case of coevolution. Two distinct pulses of network expansion predate the origin of animals, indicating that the functional diversity of human MAPK network proteins originate from ancient evolutionary radiations. Together, our study provides a critical look at the deep history of this important network.

## Introduction

The mitogen-activated protein kinase (MAPK) signaling network (Lewis et al. 1998, Cargnello and Roux 2011, Kyriakis and Avruch 2012) regulates a staggering diversity of developmental and physiological processes. For instance, both innate immune defenses against pathogens in plants and distal rib patterning in vertebrates both require MAPK signaling (Asai et al. 2002, Smith et al. 2005). In humans, MAPK pathways are implicated in diseases from cancer to COVID-19 and pose valuable therapeutic targets (Samatar and Poulikakos 2014, Weisberg et al. 2020). However, the network’s most significant role is in the governance of fundamental cellular processes as proliferation, differentiation, development, and cell survival or death (Cargnello and Roux 2011, Kyriakis and Avruch 2012). Moreover, the MAPK network is ancient and highly conserved across eukaryotic life. The most recent common ancestor (MRCA) of animals and plants is inferred to have possessed at least one functioning MAPK signaling pathway, whose origins would therefore date to nearly two billion years ago (bya; Rosindell and Harmon 2012, Betts et al. 2018).

The stereotypical MAPK pathway is a three-tier sequential phosphorylation cascade (Figure 1) in which a MAPK kinase kinase (MAP3K) transduces upstream stimuli, a MAPK kinase (MAP2K) amplifies signal, and a MAPK phosphorylates downstream effectors to ultimately modify cell behavior (Kyriakis and Avruch 2012). The human kinome includes 21-25 MAP3Ks, 7 MAP2Ks, and 14 MAPKs (Manning et al. 2002). Human MAPKs have four typical paralogs: extracellular signal-regulated kinases 1 and 2 (ERK1/2), ERK5, p38s, and c-Jun N-terminal kinases (JNKs) (Cargnello and Roux 2011, Kyriakis and Avruch 2012, Pang et al. 2022). The ERK1/2 pathway responds primarily to growth factors and is best known for regulating cell migration and cell-cycle progression. p38s and JNKs respond to stressors ranging from ultraviolet radiation, oxidative stress, osmotic stress, to inflammatory cytokines. These two “stress-activated protein kinases (SAPKs)” are often linked to cell survival or cell death. The less understood ERK5 has been linked to cell proliferation, mechanoreception, and differentiation (Paudel et al. 2021). The atypical MAPKs — ERK3/4, ERK7, and nemo-like kinase (NLK) — do not conform to the canonical three-level structure and are instead regulated by autophosphorylation or other signaling molecules (Déléris et al. 2011, Deniz et al. 2023). All MAPKs are closely related proline-directed serine-threonine kinases (Manning et al. 2002) that share a common docking (CD) domain and a kinase domain activation loop with a T-x-Y dual phosphorylation motif (Canagarajah et al. 1997, Lewis et al. 1998). Their highly conserved structure belies marked variation in nucleotide sequence, especially those coding the central ‘x’ residue of the dual phosphorylation motif, which is distinctive of specific MAPKs (Cargnello and Roux, 2011).

**Figure 1.**
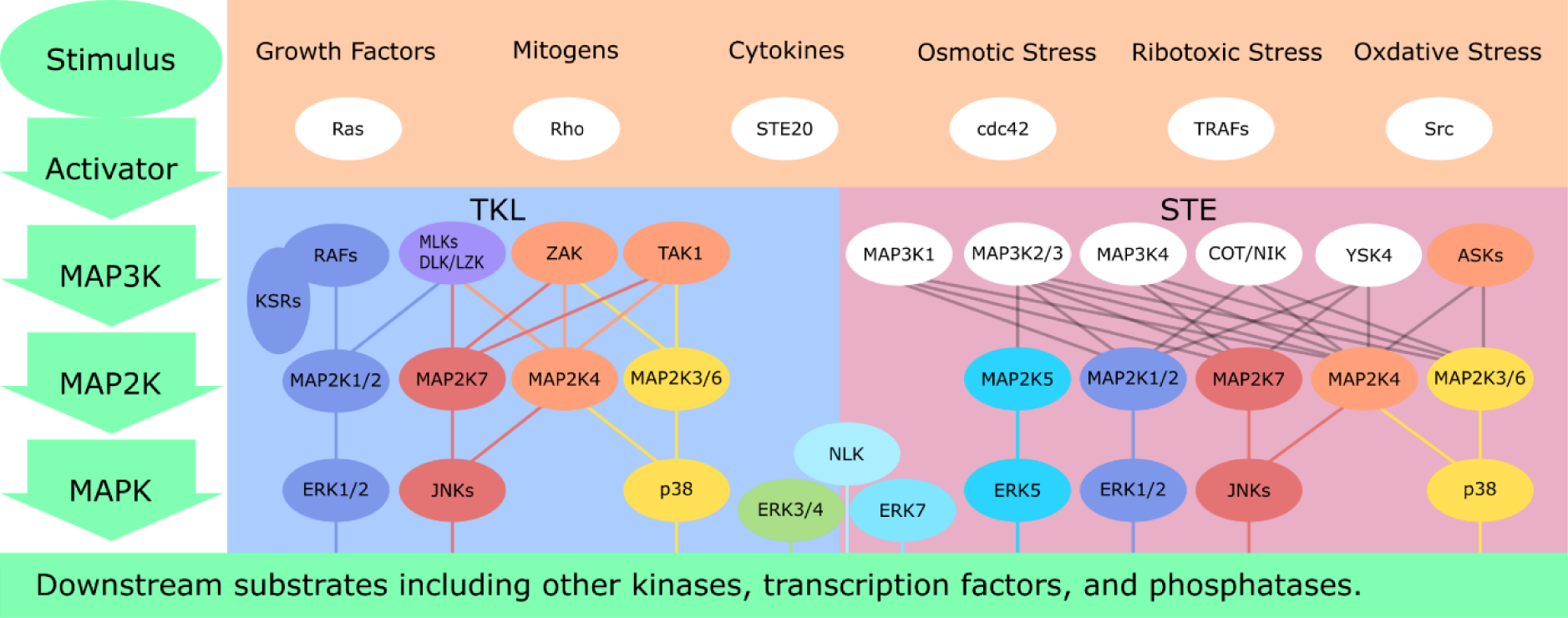
MAPK signaling network. Schematic of the three-tier MAPK signaling cascade. In a typical pathway, a MAP3K responds to upstream stimuli, including various molecules and stresses, and phosphorylates its cognate MAP2K(s). The MAP2K(s) then phosphorylate the cognate MAPK(s), which phosphorylate a variety of downstream substrates. ERK3/4, ERK7, and NLK do not have upstream MAP2K. While both TKL MAP3Ks and STE MAP3Ks phosphorylate MAP2Ks, the relationship between TKL MAP3Ks and MAP2K is clearer.

The T-x-Y motif is phosphorylated by the dual-specificity kinase MAP2Ks and is crucial for MAP2K specificity. MAPKs have well-established cognate MAP2Ks: MAP2K1/2 activates ERK1/2, and MAP2K5 activates ERK5. MAP2K3/6 activates p38s, MAP2K7 activates JNKs, and MAP2K4 activates JNKs but also p38s to a lesser extent (Cargnello and Roux 2011, Peterson et al. 2022). Like MAPKs, MAP2Ks have dual phosphorylation sites in their activation loop, but the two phospho-acceptors are positioned three residues apart (Zheng and Guan 1994). The activation loop of MAP2Ks is phosphorylated by MAP3Ks which, in addition to a kinase domain, possess large regulatory domains that connect to upstream signals or stimuli. Most recognized MAP3Ks fall into two phylogenetically distant groups: tyrosine kinase-like (TKL) MAP3Ks and sterile-like (STE) MAP3Ks (Manning et al. 2002, Peterson et al. 2022). The TKL MAP3Ks include RAFs, mixed lineage kinases (MLKs) and their relatives DLK/LZK, ZAK and TAK. The STE MAP3Ks, which belong to the same family of dual-specificity kinases as MAP2Ks, include MEKK1-4, ASKs, COT, and NIK. Recent systematic analysis of human MAP3Ks proposes that MAP3Ks govern a wide array of cell behaviors through combinatorial activation of downstream MAP2K-MAPK pathways (Peterson et al. 2022), portraying a web-like network that cannot be easily separated into independent pathways. Disentangling these complex relationships within the network remains the primary challenge to a more complete understanding of MAPK signaling.

An underexplored line of insight into MAPK network structure and function is its evolutionary history. The genetic and functional diversity observed in the human MAPK network represents approximately two billion years of evolution at all three levels of signaling (Caffrey et al. 1999, Manning et al. 2002, Li et al. 2011, Rosindell and Harmon 2012, Betts et al. 2018). Yet details of these expansions, including their genetic identity, relative and absolute timing, proximal and distal drivers, and implications for modern functionality/specificity, all remain poorly understood. Most published evolutionary studies focus solely on MAPKs (Kultz 1998, Li et al. 2011, Shabardina et al. 2023). Since the publication of the human kinome, more inclusive evolutionary studies (e.g., Goldberg 2006, Paps and Holland 2018) have been limited by a presence/absence approach that precludes a gene tree, which is necessary to identify molecular drivers of network expansion. Others are constrained by highly restricted taxon sampling and its tendency to produce unreliable tree topologies with inordinately long branches, whose utility for inferring evolutionary transformations and dynamics can be questionable (e.g., Horton et al. 2011, Li et al. 2011, Zulawski et al. 2014). The MAPK networks functionally demonstrated in various model organisms (Hadwiger and Nguyen, 2011, González-Rubio et al., 2021) suggest that at least some pathways have retained their three-tier hierarchy through evolutionary time (Dóczi et al. 2012). Hence, phylogenetic analyses that examine the multi-level network as a whole have the potential to uncover yet unappreciated evolutionary patterns.

In this study, we hypothesize that the accommodation of functionally novel paralogs would entail concomitant expansions at other levels. Our prediction therefore is that the three tiers of the MAPK network expanded in phylogenetic and functional parallel. We test this by conducting a series of phylogenetic analysis of MAP3Ks, MAP2Ks, and MAPKs across the entire breadth of the eukaryote radiation. Our curated species sample represents each of the major eukaryote radiations as well as key positions in the eukaryote tree along the human phylogenetic backbone (Strassert et al. 2021). By focusing on human paralogs, we seek to recover the position and relative timing of expansion along the direct lineage connecting the ancestral crown eukaryote to modern humans. We also infer the functions of these ancestral homologs and pathways based on reports of MAPK signaling behaviors in various extant lineages. Our objective is to provide an evolutionary perspective on the complex signaling dynamics of the modern human MAPK network.

## Materials and Methods

### Taxon Sampling

We selected a total of 42 eukaryotic species (Figure 2) representing the major lineages within Eukarya (Cantino and Queiroz 2020). Our sampling strategy was designed with the goal of inferring evolutionary events specifically along the human phylogenetic backbone. Sampled taxa also reflect the maturity of available genomic data, representing some of the most complete sets of predicted proteins for their respective eukaryotic clade on Genbank (Clark et al. 2016). Taxa whose predicted proteins were available on alternative platforms were included only when they occupied phylogenetic positions that would otherwise be poorly or un-represented in the study. These include *Ephydatia muelleri* (Kenny et al. 2020) to supplement the only other sponge (i.e., *Amphimedon queenslandica*), *Mnemiopsis leidyi* (Moreland et al. 2020) to represent Ctenophora, *Creolimax fragrantissima* (de Mendoza et al. 2015) and one *Sphaeroforma arctica* (Dudin et al. 2019) sequence to represent Ichthyosporea, *Parvularia atlantis* (Multicellgenome Lab, 2017b) to represent Nucleariida, and *Lenisia limosa* (Multicellgenome lab, 2017a) to represent Breviatea. We also attempted to sample from Asgardarchaeota, a metagenomic assemblage of archean-grade sequences which is considered to represent a sister lineage of Eukarya or a closely related paraphyletic group (Eme et al. 2023). Our analyses were unable to support any of the candidate archaean sequences as eukaryote orthologs at any MAPK level.

**Figure 2.**
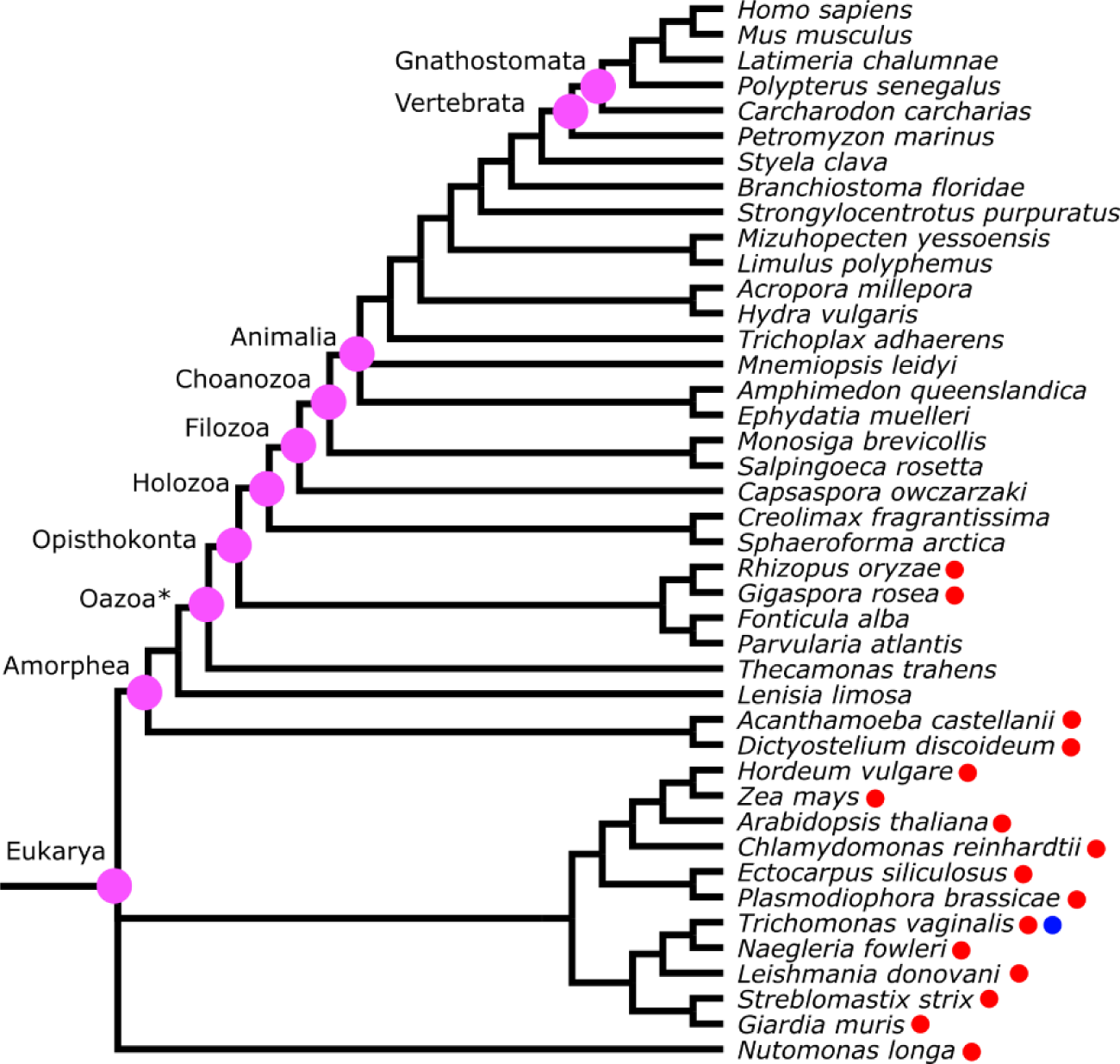
Sampled taxa. Taxa sampled in this study and their phylogenetic relationship based on OneZoom (Rosindell 2012, Betts et al. 2018). Clades mentioned in this study are labeled with their corresponding names. Taxa lacking TKL MAP3Ks are noted with red dots, and the single taxon lacking STE MAP3Ks is noted with a blue dot. Branches are not scaled to time.

### Ortholog identification

We analyzed protein sequences throughout this study because initial assessments revealed that nucleotide sequences align poorly, likely due to relatively high substitution rates outside of the conservative motifs and the broad phylogenetic scale of this study. Candidate sequences were searched for with BLAST (Johnson et al. 2008) and recorded in SI Appendix, Table S1-S4. We used blastp for all taxa that have predicted proteins in the form of amino acid sequences, and tblastn for those taxa represented only by nucleotide sequences. We began by retrieving human homologs of all three levels, as annotated on Genbank, and aligned them respectively using MAFFT version 7 (Katoh and Standley 2013) with the E-INS-I option. This produced 14 MAPKs, 7 MAP2Ks, 12 TKL MAP3Ks, and 10 STE MAP3Ks. The three TAO MAP3Ks were excluded as they are distantly related to both the TKL MAP3Ks and the STE MAP3Ks (Manning et al. 2002). KSRs, noncatalytic scaffold proteins in the RAF-MEK1/2-ERK1/2 pathway, were included for their close phylogenetic relationship with the RAFs as well as their functional relevance (Nguyen et al. 2002). Similarly, although some studies do not consider ILK and TNNI3K as MAP3Ks, they are included in this study for their apparent close phylogenetic relationship with the TKL MAP3Ks. In total, 14 TKL MAP3K paralogs were analyzed. We used TrimAl v1.3 (Capella-Gutierrez et al. 2009) with the – ‘gappyout’ option to remove poorly aligned regions for each group of kinases, resulting in four reference matrices. Sequences from other taxa were then added in an order that reflected their phylogenetic distance from humans based on the topology depicted in OneZoom (Rosindell and Harmon 2012).

We conducted an initial BLAST search for each non-human taxon with the protein sequences of ERK1, MAP2K1, MLK1, and MAP3K1, retaining the top 100 matches regardless of their e-values. These 100 candidate sequences were added to the corresponding reference matrix using the --add and --keeplength options in MAFFT. This matrix was used for maximum likelihood phylogenetic analysis with RAxML version 8.2.10 (Stamatakis 2014) with PROTCATAUTO as the protein substitution model. The resulting trees were visualized with Figtree v.1.4.4 to identify orthologs of each human paralog. At this stage, trees were rooted based on the published human kinome tree (Manning et al. 2002). Although the human kinome tree itself is unrooted, it is sufficient for rooting subtrees within each kinase family. We considered a candidate sequence to be orthologous to one or more human paralogs if: 1) it was more closely related to the specific paralog(s) than any other human paralogs, and 2) it did not cluster with the other candidate sequences to form a group that connected to the rest of the tree by a long branch. The second criterion was based on the typical behavior when distantly related sequences are included in an analysis, essentially a complex form of long-branch attraction of the outgroup (Bergsten 2005). In both MAPK and MAP2K matrices, sequences from a single reconstructed clade consisting of several non-animal opisthokonts (i.e., MRCA of fungi and animals, and all its descents) were removed due to their recovery in a phylogenetically improbable position. That position was close to the root of the entire tree, which implies that a single paralog persisted until the MRCA of opisthokonts and was subsequently lost independently in almost every major eukaryotic lineage. Long-branch attraction is the more likely scenario, with these sequences representing a unique opisthokont lineage that is not conserved in Animalia and thus outside the scope of this study. These homologs are recorded but not included in subsequent analyses.

Qualified sequences were integrated with the original matrix to produce a new reference matrix for the sequence search of the next taxon. In cases where multiple orthologous sequences exist and represent identical phylogenetic positions (either due to additional gene duplication, isoforms, or sampling from multiple, independently sequenced genomes of the same species), we randomly chose a single sequence for subsequent analysis. We made no attempts to infer evolutionary events within the various eukaryotic ‘side branches.’ To focus on events that collectively best explain the current state of the human kinome, we limited our attention to those transformations that optimize along the deep and direct history of humans. Detailing evolution within other lineages would require a significant expansion of sampled taxa and to identify all homologs within every analyzed genome—a worthy and important task, but one well beyond the scope of this study.

After going through the complete list of taxa once, we realigned and retrimmed the reference matrices with full sequences to improve alignment. We then conducted an additional round of searches, looking for orthologs that our current understanding of likely eukaryote phylogeny indicates should exist but were not identified in the initial search. For this second round, we conducted BLAST search for the taxon with ‘missing’ ortholog using the relevant homologs from closely related sampled taxa. For example, if paralog X was found in both human and sea urchin but not in lamprey, we looked for lamprey homologs using homolog X of other deuterostomes. We did not find YSK4 ortholog for the chondrichthyan *Carcharodon carcharias* but locate one for another chondrichthyan, *Scyliorhinus canicular*, and thus include this sequence instead. BLAST search results were processed as described above to further expand the matrices. Finally, to reduce Type I error (which occurs when a selected sequence belongs to another kinase group), we conducted BLAST searches for candidate sequences against the Genbank database to look for other protein families these sequences might belong to. Candidate sequences are removed if they were found likely to be orthologs of other human proteins.

### Phylogenetic Analysis and Interpretation

We selected closely related human kinases from the published human kinome tree to serve as outgroups. We then realigned and retrimmed the full sequences of all homologs along with these human kinases to create the final matrices for phylogenetic reconstruction. NLK was removed from MAPK phylogenetic analysis for two reasons: its phylogenetic instability during our reconstruction process, and the extreme length of the branch connecting the NLK sequences to the rest of the tree, which indicates substantial mutation from the original form (SI Appendix, Figure 1,2). Additional ortholog candidates of COT and NIK were added to STE MAP3K phylogeny on the basis of their scattered presence/absence pattern among non-vertebrate animals.

Phylogenetic analyses were conducted with RAxML using the PROTCATAUTO substitution model with 100 rapid bootstrap searches. Reconstructed phylogenies were rooted primarily with the outgroup kinases, and supported by the basal positions of plant homologs. Due to the scale of timespan and the limited sequence length, tree topology can be highly unstable in certain areas, and the tree space can be challenging for the algorithm to navigate. To avoid over-interpreting the divergence events, we performed a total of ten reconstruction replicates, each with the same setup except for the difference in seed numbers. We then employed the strict consensus tree derived using PAUP* 4.0a169 (Swofford 2003) for subsequent interpretations, and consider the reconstruction with highest likelihood among the ten replicates when confirming finer inferences (SI Appendix, Figure S3-S10).

Divergence time of protein paralogs were subsequently determined based on topology and taxonomy around nodes that indicate separation of the human homologs. Ancestral states and orthologs were inferred according to the most parsimonious interpretation of amino acid substitution. For example, the dual phosphorylation motif of MRCA of all JNK homologs (*_MRCA_*JNK) is reconstructed as T-P-Y, based on the conserved T-P-Y pattern in all JNK homologs on the MAPK phylogeny. No numerical divergence time estimations were made because several branches represent millions if not more than a billion years of evolution, and any node estimates would be unreliable for the sampling density in this study at this time scale.

### Phosphorylation motif

The residues immediately flanking substrate phosphorylation sites play a critical role in kinase specificity (Ubersax and Ferrell Jr 2007). We examined the highly conserved dual phosphorylation motif in the activation loops of MAP2Ks and MAPKs as means of tracing functional shifts. In particular, we focused on differences in charge and hydrophobicity, which are often indicative of shifts in binding preference (Ubersax and Ferrell Jr 2007). For MAPKs, we examined the variable central residue between the dual phosphorylation sites of the signature T-x-Y motif (Cargnello and Roux 2011). In MAP2Ks, the dual phosphorylation sites are variably separated by three to five residues, and the motif is less conserved. To capture the residues most likely to affect specificity, we examined from the residue immediately preceding (p-1) the first phosphorylation site to the residue immediately following (p+1) the second phosphorylation site.

## Results

### MAPK Phylogeny

Our search identified at least one MAPK homolog in all 42 examined eukaryotes (Figure 2). Our phylogeny points to ERK7 as the earliest MAPK to diverge from the rest of the MAPKs (Figure 3,4). We found ERK7 orthologs to be distributed across the span of eukaryotic diversity (SI Appendix, Table S1), which suggests that the formative divergence of ERK7 occurred prior to the approximately two billion-year-old MRCA of eukaryotes. We infer that the ancestral MAPK possessed the T-E-Y dual phosphorylation motif in its activation loop. This primitive motif is maintained in ERK1/2, ERK5, and ERK7 with few exceptions.

**Figure 3.**
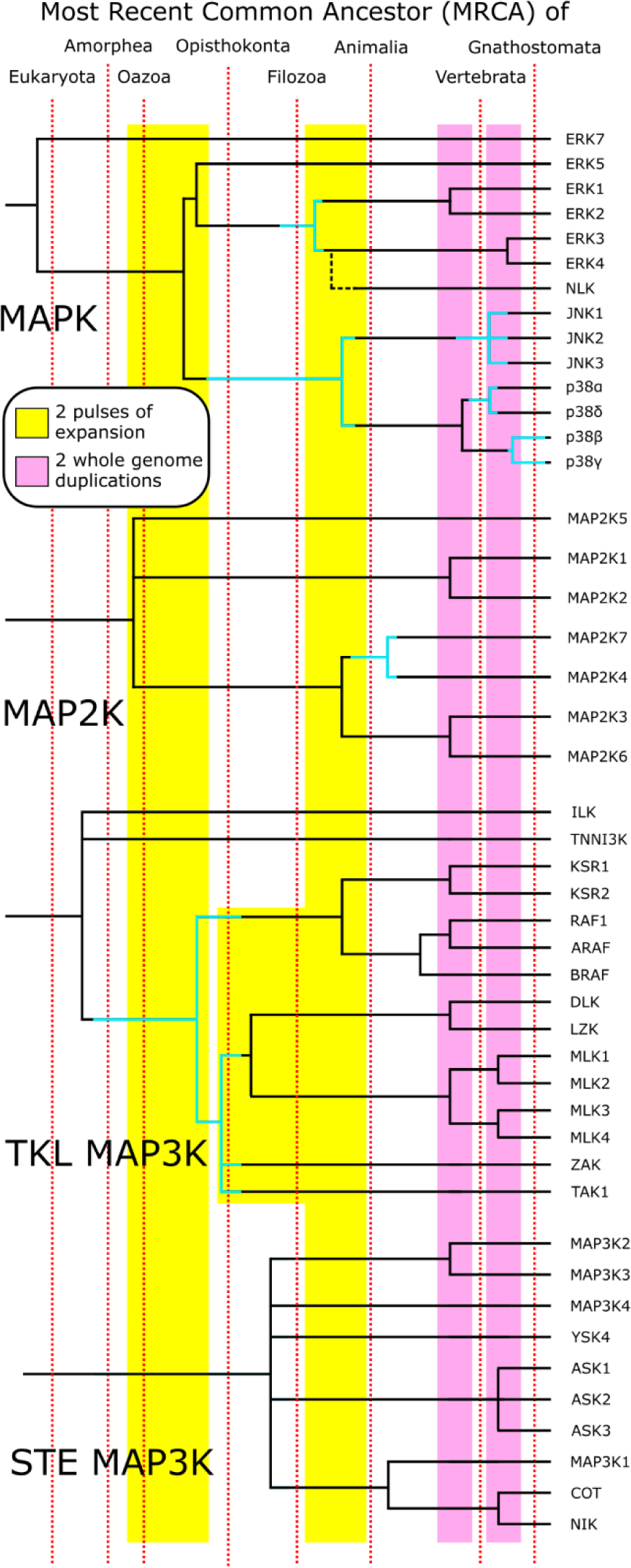
Phylogenies of the MAPK signaling network. Phylogenies of human MAPK, MAP2K, TKL MAP3K, and STE MAP3K. Each tree summarize ten replicates of maximum likelihood estimations under a strict consensus criterion. An exception is made at the origin of TKL MAP3K: the concensus tree depict a four-way polytomy, but the divergence of ILK and TNNI3K from the TKL MAP3Ks is supported by 8 out of 10 replicates along with their functional distinctiveness. Their functional distinctiveness further supports their depiction in this figure. Divergence time estimates are based on taxonomic representation of the homologs surrounding the respective nodes. Blue branches represent divergence events whose time of occurrence are ambiguous due to possible loss of homologs, with the interval representing possible timing of divergence. The two pulses of expansion and the two WGDs are highlighted. Trees are not scaled to time.

**Figure 4.**
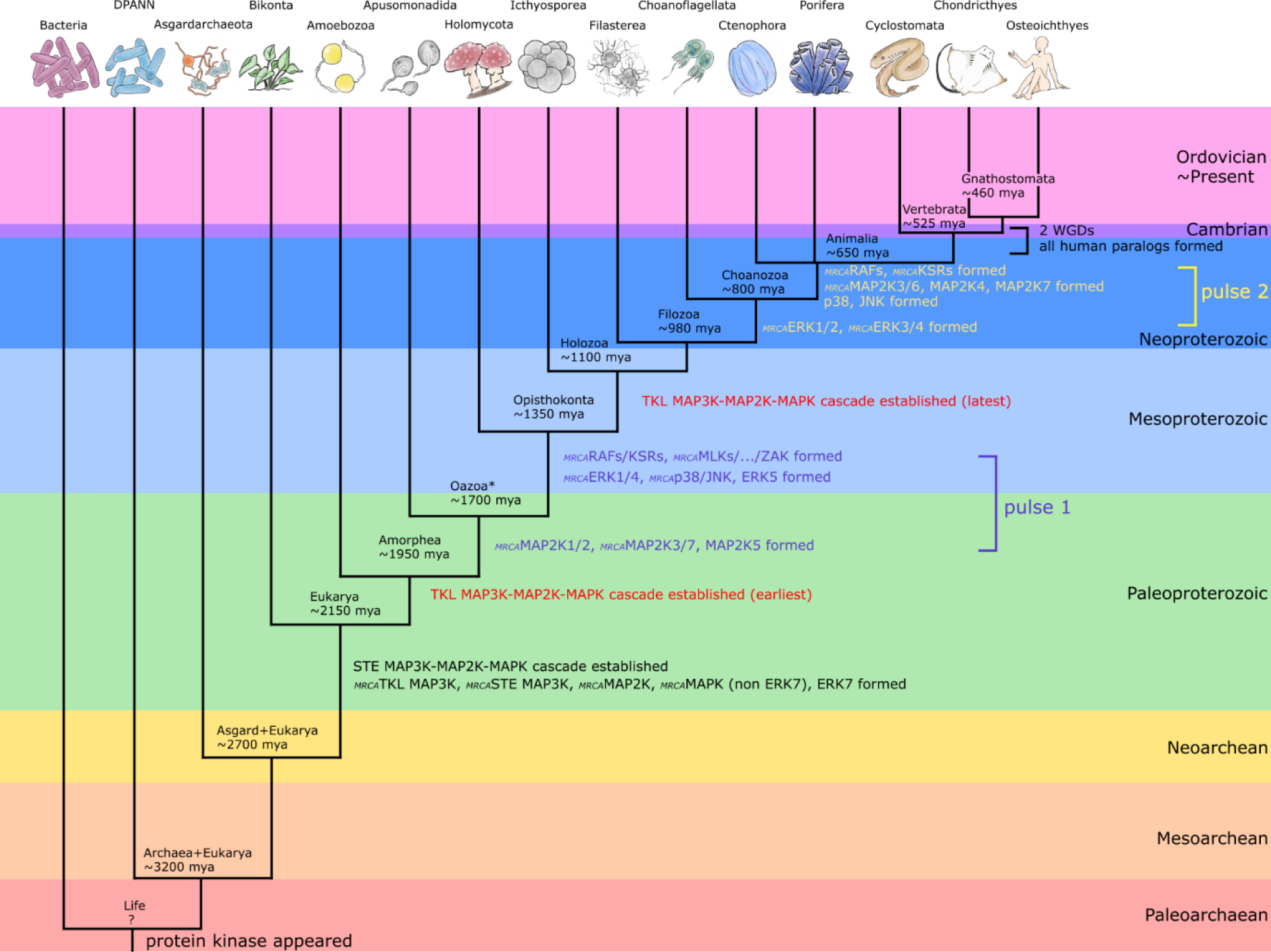
Evolutionary history of the MAPK signaling network. Key events of the MAPK signaling network evolution mapped onto the tree of life (scaled to time), with a focus on the two pulses of expansion. Both pulses involve TKL MAP3K, MAP2K, and MAPK, no evidence of STE MAP3K involvement is found. The first MAPK signaling pathway likely appeared on the stem of Eukarya, and all paralogs found in humans were formed prior to the MRCA of gnathostomes. Divergence time estimates of taxa are based on OneZoom (Rosindell 2012, Betts et al. 2018), whereas divergence time of paralogs are inferred from phylogenies reconstructed in this study.

We optimized the first pulse of canonical MAPK expansion to the stem of Opisthokonta (1.7-1.35 bya), where a single ancestral MAPK became three functionally distinct paralogs. The yeast *Saccharomyces cerevisiae* has three MAPK paralogs Fus3/Kss1, Hog1, and Slt2 (Lewis et al. 1998, Bardwell 2006). Our phylogeny shows that Fus3/Kss1 is the ortholog of *_MRCA_*ERK1/4 (i.e. most recent common ancestral protein of ERK1 and ERK4), Hog1 is the ortholog of *_MRCA_*p38/JNK, and Slt2 is the ortholog of ERK5. These three kinase lineages were found in all sampled opisthokonts except for a few species that potentially lack ERK5, including the horseshoe crab *Limulus polyphemus* and the tunicate *Styela clava*. At the protein sequence level, the vast majority of sampled MAPKs retain the plesiomorphic form of the dual phosphorylation motif, T-E-Y. *_MRCA_*p38/JNK notably deviates from this pattern by replacing the central glutamate with glycine. An exception is the holomycotan *Fonticula alba*, whose *_MRCA_*p38/JNK ortholog retains the glutamate. A few taxa, such as the sea urchin *Strongylocentrotus purpuratus* and the choanoflagellate *Salpingoeca rosetta*, also have a substituted middle residue in ERK5.

ERK3/4, a lineage of atypical MAPKs, was previously recovered as a basal divergence among MAPKs (Manning et al. 2002). Our trees however indicate that *_MRCA_*ERK1/4 next produced a distinct *_MRCA_*ERK1/2 and *_MRCA_*ERK3/4 along the stem lineage of Holozoa (1.35-1.1 bya). Orthologs of *_MRCA_*ERK1/2 but not *_MRCA_*ERK3/4 were inferred in Icthyosporea and Filasterea. In contrast, orthologs of *_MRCA_*ERK3/4 can only be inferred for Choanozoa onwards, raising the possibility that *_MRCA_*ERK3/4 did not diverge from *_MRCA_*ERK1/2 until the choanozoan stem lineage (980-800 mya).

Our analysis confirms that the evolutionary bifurcation of *_MRCA_*p38/JNK into p38 and JNK happened either prior to or on the stem of animals (800-650 mya). The strict consensus tree recovered JNK orthologs for every animal included in this study, whereas p38 and *_MRCA_*p38/JNK (i.e., hog1) orthologs form a large polytomy along with *_MRCA_*JNKs. The reconstruction with highest likelihood recovered at most one homolog from all but one non-animal taxa as orthologs of *_MRCA_*JNK, *_MRCA_*p38, or *_MRCA_*JNK/p38, *Monosiga brevicollis*, one of the two choanoflagellate included in this study, possesses inferred ortholog for both p38 and JNK. The separation of p38s and JNKs marks the final divergence of any MAPK in the human phylogenetic backbone that is associated with significant functional innovation.

The stem lineage of vertebrates (535-525 mya) and gnathostome (525-460 mya) later witnessed another expansion that resulted in seven additional MAPK paralogs, which are likely the product of two whole genome duplications (WGDs) that are well-established for this part of the tree (Dehal et al. 2005). Similar expansions are observed at other levels of the signaling network. Despite increasing the genetic diversity on all tiers, the two WGDs in early vertebrate evolution (Nakatani et al. 2021) produced only functionally similar and potentially redundant paralogs (Cargnello and Roux 2011). As noted previously, WGDs are not always associated with marked increases in functional diversity (Van de Peer et al. 2009).

### MAP2K Phylogeny

At least one MAP2K homolog was identified in all 42 sampled eukaryotes (Figure 2). Our reconstruction (Figure 3,4) places the first pulse of MAP2K diversification along the stem of Oazoa (1.8-1.7 bya), a group we define as including all Obazoan taxa except the most basal Breviatea (Brown et al. 2013). This expansion produced a triad of MAP2K lineages, MAP2K5, *_MRCA_*MAP2K1/2 and *_MRCA_*MAP2K3/4/6/7. Their collective origin predates that of their downstream targets ERK5, ERK1/2, and *_MRCA_*p38/JNK. We recovered no clear orthologs of MAP2K5 in Apusomonadida and Fungi. These species do possess sequences we tentatively interpret as MAP2K5 orthologs based on their recovered position near the *_MRCA_*MAP2K5 and as well as functional similarities between human MAP2K5 and the inferred yeast orthologs (Jo et al. 2017). This initial diversification was followed by a considerable period of MAP2K conservation, during which MAP2K5 and MAP2K1/2 exhibit no further splitting events.

In contrast, *_MRCA_*MAP2K3/6 and *_MRCA_*MAP2K4/7 diverge along the stem of Animalia, paralleling the emergence of their targets, the p38s and JNKs. One homolog each for the two sampled sponges and two ctenophore homologs were recovered basal to *_MRCA_*MAP2K3/4/6/7. We interpret the two ctenophore sequences as orthologs of *_MRCA_*MAP2K3/6 and MAP2K4. As clear orthologs for other MAP2K lineages are present, we interpret the sponge sequences as orthologs of MAP2K4. A final MAP2K expansion on the vertebrate stem lineage marks the splitting of MAP2K1 and MAP2K2, and of MAP2K3 and MAP2K6. Like their cognate MAPKs, these divergences likely correspond to vertebrate WGDs and are not associated with clear functional novelty.

### TKL MAP3K Phylogeny

The MAP3K phylogenies of both TKLs and STEs lack the level of resolution attained in the MAPK and MAP2K analyses (Figure 3). TNNI3K and ILK, two TKLs that uniquely share an ankyrin repeats domain (ANK) (Lal et al. 2014), diverged from the TKL MAP3Ks on the stem of Amorphea. This initial divergence was followed by the splitting of *_MRCA_*RAF/KSR from the MRCA of all remaining TKL MAPKs (Figure 4). This event is inferred to the stem of Opisthokonta and evidenced by the observation that several non-fungi holomycotans possess orthologs of specific TKL paralog within the clade defined by the divergence of ZAK and MLKs. The subsequent split of the RAFs and KSRs happened on the stem of Animalia. Precise mapping of TAK1, ZAK, and *_MRCA_*DLK/MLK divergence is difficult due to multiple inferred losses in non-animal taxa, but it must have occurred no later than the origin of crown Opisthokonta. The later expansions of the paralogs of RAF, KSR, and MLK, as well as the separation of DLK and LZK, all appear related to the WGDs that mark the origin of vertebrates.

TKL MAP3Ks do not show the level of evolutionary conservation within Eukarya that marks both MAPKs and MAP2Ks, with inferred losses in several early diverging lineages (Figure 2). For example, plants possess only the TNNI3K orthologs, but no orthologs of TKL MAP3Ks. In addition, Fungi appear to have lost all TKL MAP3Ks, whereas the closely related Nucleariida retains only orthologs of MLKs and DLK/LZK. Ichthyosporea has only one TKL MAP3K sequence that corresponds to TAK1. This lineage does conserve a clear ortholog of *_MRCA_*RAF/KSR, which neither Fungi nor Nucleariida possess. The collective distribution of TKL MAP3K orthologs is thus complex, rendering the divergence-time estimation a challenge. This is especially the case for events predating the origin of Animalia.

### STE MAP3K Phylogeny

At least one STE MAP3K sequence was identified in each of the sampled taxa except in *Trichomonas vaginalis* (Figure 2). The phylogeny of STE MAP3Ks is limited by the low resolution at the base of the radiation, extreme branch lengths, and some nested relationships that seem highly unlikely (Figure 3). For example, a sequence from the early diverging eukaryote *Leishmania donovani* was reconstructed as the sister homolog of a sequence from the choanoflagellate S*alpingoeca rosetta*. In addition, there are multiple inferred paralog losses even within Animalia; the sampled sponges notably lack a MAP3K1. The technical replicates together produce five different tree topologies even just considering the human paralogs, emphasizing the ambiguity concerning the early divergence of STE MAP3Ks. Such complicatizons make rendering the details of STE MAP3K phylogeny a challenge. Nevertheless, all but two Obazoans have at least one sequence that bears strong resemblance to a specific human STE MAP3K, suggesting that there is still some extent of resolution.

The strict consensus tree portrays the origin of STE MAP3Ks as a large polytomy, but resolution increases as it gets closer to the origin of animals. Within Animalia, there are clearly established lineages: *_MRCA_*MAP3K1/COT/NIK, *_MRCA_*MAP3K2/3, MAP3K4, ASKs, and YSK4. These five lineages were definitely established prior to crown Animalia, possibly as early as our split from the amoebozoans. The final pulse of expansion again relates to the two vertebrate WGDs, where MAP3K2 and MAP3K3, COT and NIK, and the ASKs paralogs diverged from one another.

## Discussion

### TKL MAP3Ks, STE MAP3Ks, and the origin of the MAPK signaling network

We find STE MAP3Ks, MAP2Ks, and MAPK orthologs to be universally present across all sampled lineages with a single exception – *Trichomonas vaginalis*. Whether such an exception is due to incomplete data or a true loss remains unclear. Plants, our most distant eukaryotic relatives, have STE MAP3Ks that are shown experimentally to function in classical three-tier phosphorylation cascades with MAP2Ks and MAPKs (Zhang et al. 2022). In diverse taxa, STE MAP3Ks have been shown experimentally to function as MAP3Ks at the top of a classic three-tier MAPK cascade, by phosphorylating and activating a downstream MAP2K (e.g., Herskowitz 1995, Jin et al. 2002, Guo et al. 2021). Together, this provides robust phylogenetic evidence that the basic three-tier architecture of the MAPK network was established and functional prior to the earliest divergence of eukaryotes, and that the original pathway likely was regulated by a STE MAP3K.

STE MAP3Ks across Eukarya respond to external stimuli, especially environmental stressors such as osmotic stress, oxidative stress, UV radiation, and infection or inflammation (Duch et al. 2012, Peterson et al. 2022). In response to STE MAP3K activation, the downstream MAP2K-MAPK pathways induce pro-survival adaptations including cell cycle regulation and differentiation. Their ability to coordinate cell fate decisions is also harnessed for processes such as proliferation, migration, development, or even programmed cell death. In yeast, MAP3Ks (which are all STEs) mediate not only adaptive responses to osmotic stress (de Nadal and Posas 2022) but also sexual reproduction (Herskowitz 1995) and differentiation (Roberts and Fink 1994). Three-tier STE MAP3K pathways in plants regulate biotic and abiotic stress responses as well as tissue development (Jin et al. 2002). In the dinoflagellate *Scrippsiella trochoidea*, cold and darkness-triggered encystment induces upregulation of genes related to MAPK, MAP2K, and a STE MAP3K (Guo et al. 2021). Overall, phylogenetic bracketing indicates that the ancestral MAP2K-MAPK pathway controlled how the ancient eukaryotes altered their behavior in response to extracellular stimuli sensed by STE MAP3Ks, from stressors to developmental cues. The deepest history of STE MAP3Ks is difficult to decipher, but it is clear that the STE MAP3K-MAP2K-MAPK cascade is a buffered network that likely maintained its role in stress response for billions of years. It appears that the STEs largely maintained their role in stress as well as their ability to signal to all downstream pathways. Indeed, recent work showed that the STE MAP3Ks MAP3K1-4 are the only MAP3Ks able to simultaneously activate all MAP2Ks in human cells (Peterson et al. 2022). STE MAP3Ks with narrower affinities, such as COT/NIK, YSK4, and ASKs, are perhaps better interpreted as products of apomorphic functional reductions or losses.

In contrast, TKL MAP3Ks are not omnipresent within Eukarya. TKLs are conspicuously missing in Fungi, even though they are preserved in lineages that are further distant from humans, as well as in their holomycotan relatives. Ambiguous positions of non-Opisthokonta homologs currently impedes the precise phylogenetic placement of the initial TKL MAP3K expansion. There are also inherent limitations in ordering events that phylogenetically optimize to the same stem lineage. In addition, to date, there is no experimental evidence in the literature for TKL MAP3K orthologs participating in a classical MAPK cascade within early diverging lineages, such as plants and holomycotans. Instead, plants “Rafs” such as *Arabidopsis* CTR1 regulate targets by other mechanisms including inhibitory phosphorylation of transcription factors (Park et al. 2023). We would also like to point out that based on our phylogenetic analysis, these Raf-like kinases are not orthologs of Rafs, but rather of *_MRCA_*TLK MAP3K as a whole. In other words, the MRCA of Eukarya possess neither a TKL kinase that is nested within the clade defined by the human TKL MAP3Ks nor one that function as a MAP3K, and that TKL MAP3Ks were recruited to a previously established MAPK signaling network later in history. Indeed, our analysis places STE MAP3K-MAP2K-MAPK signaling at the origin of Eukarya where it likely predates the TKL MAP3K-MAP2K-MAPK signaling by hundreds of millions of years.

While we identify at least one ortholog for each tier of the MAP3K-MAP2K-MAPK network in nearly all sampled eukaryotic taxa, definitive MAPK network orthologs are currently unknown in Bacteria or Archaea. We were also unable to identify MAPKs in Asgardarchaeota, which are thought to be either a monophyletic sister taxon to, or a paraphyletic group inclusive of Eukarya (Zaremba-Niedzwiedzka et al. 2017). The inferred presence of a MAPK signaling pathway in MRCA of eukaryotes and the absence of archaean homologs at any level of the network together places the origin of the first MAPK pathway along the stem lineage of Eukarya, a clade apomorphically defined by the presence of a cell nucleus (Cantino and Queiroz 2020). Nonetheless at this time, Asgardarchaeota remains known only through metagenomic assembly. It is thus premature to assume any evolutionary relationship between the emergence of the eukaryotic nucleus and the MAPK pathway. Whether components of the MAPK signaling network will ultimately be identified in non-eukaryotic relatives such as Asgardarchaeota is an interesting question for future research.

### Expansion of the MAPK signaling network

The ancestral eukaryotes possessed an autoactivating ERK7 and an ancestral canonical MAPK that was activated by a single ancestral MAP2K and MAP3K. Approximately 450 million years after the MRCA of eukaryotes, all three levels of the MAPK network expand in parallel (Figure 5). On the stem of Oazoa, the ancestral MAP2K differentiated into three distinct paralogs: *_MRCA_*MAP2K1/2, *_MRCA_*MAP2K3/4/6/7, and MAP2K5. The ancestral MAPK followed suit on the stem of Opisthokonta, producing *_MRCA_*ERK1/2, *_MRCA_*p38/JNK, and *_MRCA_*ERK5. The close evolutionary and functional association between MAPK and MAP2K is suggestive of a coevolutionary relationship (Fraser et al. 2004). Meanwhile, both TKL and STE MAP3Ks experienced their own expansion. By the origin of crown Opisthokonta, *_MRCA_*RAF/KSR, now consists of exclusive MAP2K1/2-ERK1/2 regulators, diverged from *_MRCA_*ZAK/MLK. The divergence of MAP3K2/3, the only known MAP3K that has the additional ability to activate MAP2K5-ERK5, from the other STE MAP3Ks might have also happened around this period, although the timings is rather ambiguous, ranging from the origin of crown Amorphea to the origin of crown Filozoa.

**Figure 5.**
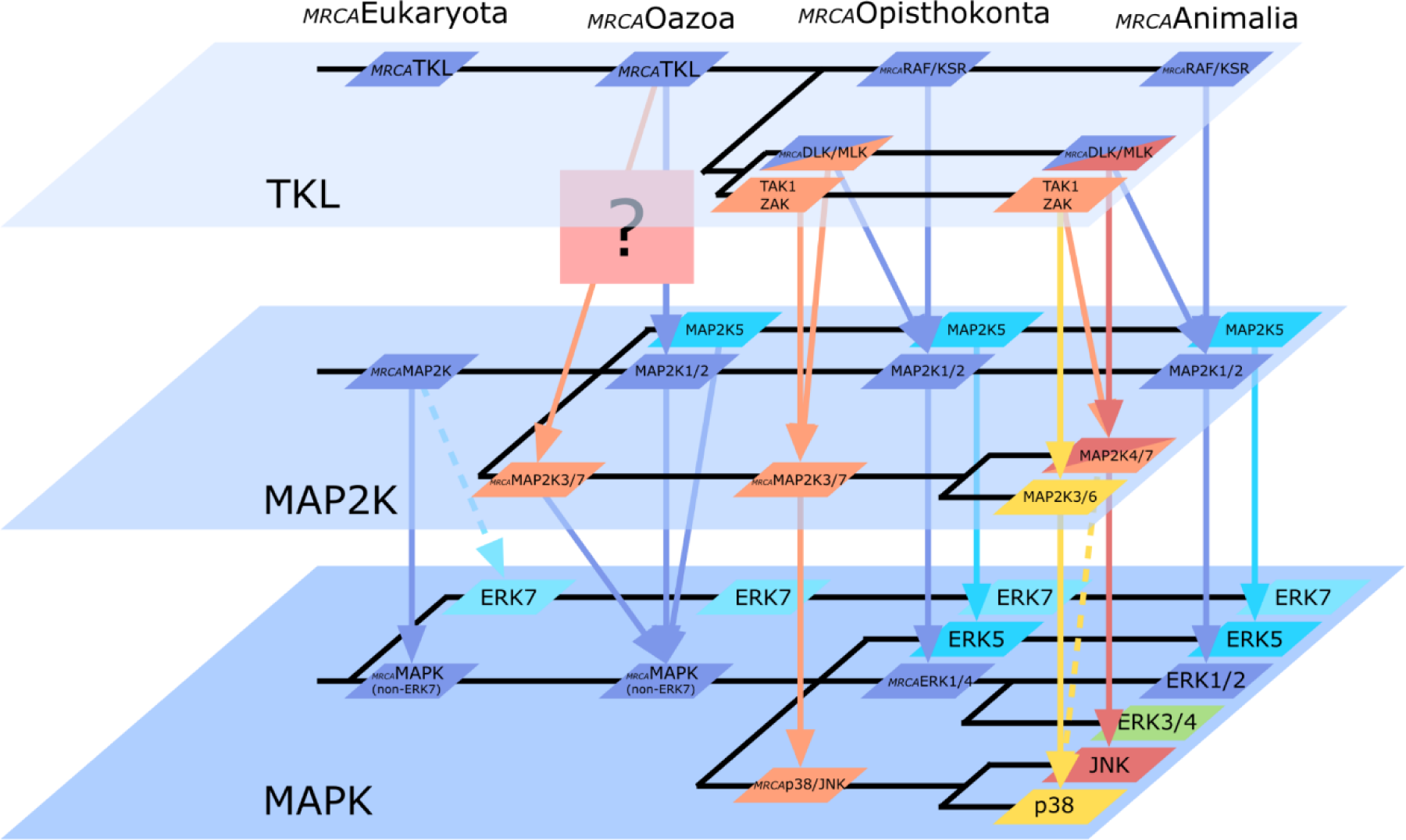
Evolutionary and functional expansion of the TKL-MAP2K-MAPK signaling cascade. A schematic showing the incorporation of TKL MAP3Ks into MAPK signaling, and the two-pulse expansion of the TKL-MAP2K-MAPK network. Function and specificity are based on functional data from extant organisms, and then inferred with ancestral state reconstruction using parsimony criterion. Lack of functional data for *Thecamonas trahens* post a challenge to infer whether TKL MAP3K ortholog in MRCA of Oazoa functions as a MAP3K. Colors of the kinases and arrows indicate general downstream specificity.

Based on the expansion described above, we conjecture that no later than the MRCA of Opisthokonta, three functionally distinct MAPK pathways were established: *_MRCA_*RAF/KSR→ *_MRCA_*MAP2K1/2→ *_MRCA_*ERK1/2, involved in proliferation and cell cycle control; *_MRCA_*ZAK/MLK→ *_MRCA_*MAP2K3/4/6/7→ *_MRCA_*p38/JNK, associated with stress response and cell fate/survival; STE MAP3K2/3 (which has a general affinity)→ MAP2K5→ ERK5, involved in mechanoreception, proliferation, and differentiation (Drew et al. 2012, Nithianandarajah-Jones et al. 2012, Tusa et al. 2018, Pereira et al. 2020). These pathways appear to have largely retained their respective functions throughout evolution, alongside the *_MRCA_*STE MAP3K’s general affinity for *_MRCA_*MAP2K. In terms of human evolution, this pulse dates to the period after the animal lineage’s divergence from Amoebozoa but predates our divergence from fungi.

The second pulse of multi-level expansion occurs on the *_MRCA_*ZAK/MLK-*_MRCA_*MAP2K3/4/6/7-*_MRCA_*p38/JNK pathway. This time, distinct paralogs at the MAP3K level appear first, significantly predating diversification of the lower levels. In our analysis, MLKs, LZK/DLK, TAK1, and ZAK became independent TKL lineages no later than crown Opisthokonta. Only much later at stem Animalia (800-650 mya), a functionally distinct/novel JNK diverges from p38. The division of their MAP2K partners optimizes to the same period, resulting in two new pathways, *_MRCA_*MAP2K3/6 – p38 and *_MRCA_*MAP2K4/7–JNK. Interestingly, parallel expansion is again more tightly coordinated between MAP2Ks and MAPKs relative to the upstream MAP3Ks.

Our findings suggest that functionally distinct homologs at one level was paralleled by mutations at the other levels to form a novel three-tier pathway. Specifically, many of the events, including the two pulses, fit an evolution model where a higher-level expansion is followed by a lower-level expansion. Based on our results, we propose that the diversification of TKL MAP3Ks was a key driver of tandem expansion of pathways across the MAPK network. Despite their late recruitment, the tandem phylogenetic and functional expansions of MAPKs and MAP2Ks along the human lineage occur in lockstep with TKL MAP3Ks.

The phylogeny of STE MAP3Ks does not show this parallelism with downstream targets, even when the basal polytomy is resolved to maximize congruency with reported functions. However, our analysis also finds it as the only component of the MAPK signaling network that witnessed a functional innovation within Animalia. The human kinome tree previously mapped COT and NIK to a basal position among STE MAP3Ks (Manning et al. 2002), but our analysis reconstructs these STEs as sister homologs to MAP3K1. COT and NIK both function as key regulators of inflammatory cytokine signaling in the immune response (Thu and Richmond 2010, Gantke et al. 2011, Buchmann 2014). Adaptive immunity, which relies on extensive cytokine networks to coordinate immune cells, is widely accepted to be a relatively derived animal trait that arose in vertebrates prior to the divergence of jawed and jawless fish (Flajnik and Kasahara 2010). While an exhaustive search revealed potential orthologs in some non-vertebrate taxa, unambiguous orthologs of COT and NIK exist only within Vertebrata. This discovery moves the origin of these two MAP3Ks to a much later date and into a more nested position among the STE radiation, likely after the origin of animals.

### Evolution by specialization

Many STE MAP3Ks have broad affinity for almost all MAP2Ks. On the other hand, TKL MAP3Ks often show overlapping but specific affinities for two or more downstream pathways. For example, TAK1 and ZAK activate the pathways MAP2K3/6-p38 and MAP2K4/7-JNK. MLK and DLK activate both MAP2K1/2-ERK and MAP2K4/7-JNK (Robitaille et al. 2010, Peterson et al. 2022). Our phylogeny suggests a parsimonious explanation for this complex pattern amongst TKL MAP3Ks: specialization via the gradual exclusion of binding partners. We propose that the primary driver of the MAPK signaling network expansion was the gradual narrowing and finetuning of TKL MAP3K specificities.

From our reconstructed phylogeny, we infer that the primordial *_MRCA_*TKL MAP3K could broadly activate *_MRCA_*MAP2K and its descendants. We hypothesize that subsequent pathway divergences reflect the gradual exclusion of binding partners. According to this scenario, the early divergence of *_MRCA_*RAF/KSR from *_MRCA_*ZAK/MLK produced two lineages with different specificity: *_MRCA_*RAF/KSR is only capable of interacting with MAP2K1/2, whereas *_MRCA_*ZAK/MLK is inferred to have retained the ancestral affinity for all MAP2Ks. As the *_MRCA_*ZAK/MLK lineage further diversified, we propose that each branch lost affinity for certain downstream pathways. Coevolution of MAP2Ks that repel specific upstream MAP3Ks would further reinforce pathway diversification. TAK1 and ZAK lost affinity for MAP2K1/2 (Peterson et al. 2022). The MLK/DLK lineage lost affinity for MAP2K3/6. When DLK and LZK later diverged in the vertebrate WGDs, LZK additionally lost affinity for MAP2K1/2 (Ikeda et al. 2001).

We also propose that the same process of expansion by exclusion was echoed downstream at the MAP2K-MAPK level, resulting in narrower and more specialized pathways. This hypothesis of gradual exclusivity is supported by the preferences of human MAP2Ks. As mentioned previously, human MAP2K4 activates both JNK and p38 (albeit with less efficacy) whereas MAP2K7 is exclusive for JNK (Cargnello and Roux 2011, Peterson et al. 2022), from which we infer that the ancestral MAP2K4/7 could activate both JNK and p38. Meanwhile, all human MAP2K3/6 paralogs exclusively activate p38 (Cargnello and Roux 2011). We can thus infer that the common ancestor of MAP2K3/6 and MAP2K4/7 also activated both p38 and JNK, and that this more plesiomorphic form and function is preserved in modern human MAP2K4. Meanwhile, MAP2K3/6-JNK and MAP2K7-p38 interactions were lost.

Key variations in primary structure of MAPKs also support this hypothesis. All human p38s and non-human p38 orthologs in the *_MRCA_*p38/JNK lineage are characterized by a T-G-Y motif, where the negatively charged glutamate of the ancestral T-E-Y motif as preserved in other MAPKs has been replaced with apolar glycine. The glycine in *_MRCA_*p38/JNK may have disfavored interactions with MAP2K1/2 or MAP2K5, which would expect a charged residue on the substrate binding site. In taxa that possess only one homolog of the *_MRCA_*p38/JNK lineage, its motif reads T-G-Y. In contrast, human JNKs and non-human JNK orthologs have T-P-Y. Both glycine and proline are amino acids whose side groups are critical determinants of protein structure (Krieger et al. 2005). However, the JNK T-P-Y motif is particularly unusual: most non-proline-directed serine/threonine kinases (such as MAP2Ks) disfavor substrates with proline at the p+1 position as the cyclic structure of proline precludes backbone hydrogen bonding (Ubersax and Ferrell Jr, 2007). We speculate that the proline may have functioned to repel other MAP2Ks and discourage non-cognate interactions.

Evolution of the cognate MAP2K active site towards accommodating the proline ring (usually with an apolar pocket) may have had the cost of reducing compatibility with glycine, which does not have a side chain. Indeed, MAP2K7 uniquely possessed a hydrophobic pocket unseen in other MAP2Ks (Murakawa et al. 2020). This may explain why MAP2K7 is specific for only JNK while MAP2K4 has broader affinity for both p38 and JNK.

Intriguingly, we observe that other kinases in the JNK pathway have unusual activation loop motifs where dual phosphorylation by upstream partners occur. Orthologs of the JNK cognate MAP2K4/7 also have a distinctive activation loop that sets them apart from other MAP2Ks (Figure 6A). Each of the two phosphorylation sites is immediately followed by a positively charged amino acid, either an arginine or a lysine. This level of conservation suggests that the positive charges at p+1 may be of functional significance. Although the MAP2K4 of some taxa lacks this residue, the second p+1 position is predominantly occupied by arginine in the majority of MAP2K4/7 orthologs. In fact, nearly all MAP2K7s conserve the motif S-K-A-x-T-R, where the second p+1 arginine is accompanied by a first p+1 lysine. Overall, these structural shifts support the idea that the MAP2K7-JNK signaling pathway represents subsequent specialization of a more inclusive ancestral pathway rather than acquisition of function de novo.

**Figure 6.**
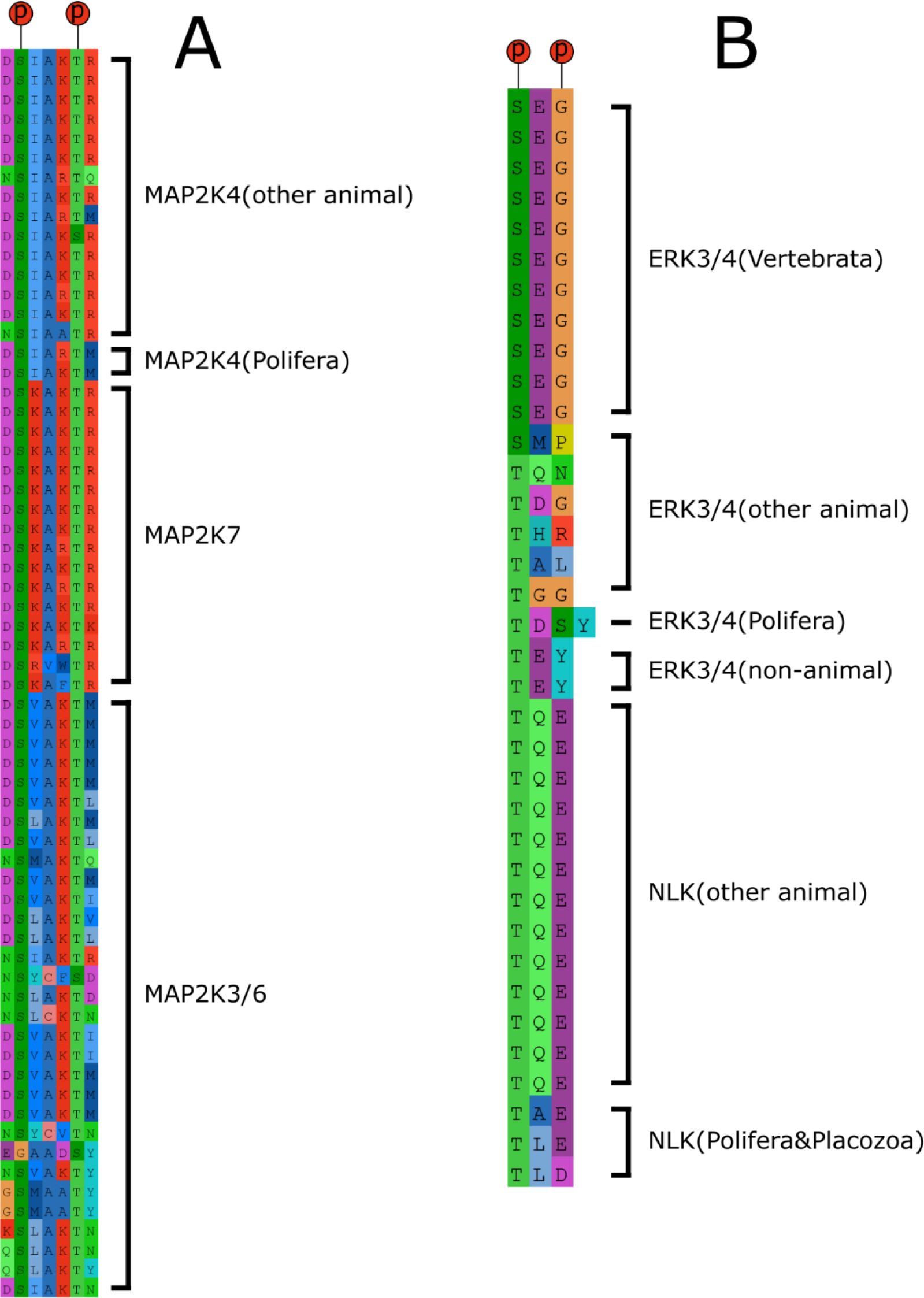
Phosphorylation motif of selected MAPKs and MAP2Ks. A) Phosphorylation motifs of the *_MRCA_*ERK3/4 and NLK orthologs. B) Phosphorylation motifs of the *_MRCA_*MAP2K3/6, MAP2K4, and MAP2K7 orthologs. The ‘p’s in red circles indicate sites homologous to phosphorylation sites in the ancestral MAPK and MAP2K. Amino acids are abbreviated using standard abbreviations. Only Threonine (T), Serine (S), and Tyrosine (Y) are capable of phosphorylation. The phosphorylation motif is highly conserved that there is no insertion or deletion observed, with the single exception being *_MRCA_*ERK3/4 ortholog in sponge *Amphimedon queenslandica*.

Our hypothesis that expansion is a result of specification and sub-functionalization rather than neo-functionalization can resolve questions such as the relationship between p38, JNK, and the yeast homolog Hog1. Hog1 is often considered equivalent to p38 (Takekawa et al. 1997) on the basis of high sequence similarity and shared roles in stress response. However, sustained activation of Hog1 can reportedly induce cell death, behavior not seen with p38 and more reminiscent of the pro-apoptotic JNK (Maeda et al. 1994, Bridge et al. 2010, You et al. 2013, Peterson et al. 2022). Experimentally, either mammalian p38 or JNK can interchangeably rescue the phenotype of Hog1-deficient *Saccharomyces cerevisiae* (Böhm et al. 2002). The functionality of Hog1 in non-animal taxa may thus be better described as a combination of the roles of animal p38 and JNK. In our data, all sampled animal genomes contain distinct p38 and JNK orthologs, whereas all sampled non-animal opisthokonts have orthologs of Hog1. We propose that Hog1 conserves the plesiomorphic state of *_MRCA_*p38/JNK, and that the emergence of distinctive p38 and JNK isoforms represents a division of labor from the ancestral all-purpose homolog.

### Atypical MAPKs

In our phylogeny, the first branch to diverge from the ancestral MAPK is the atypical ERK7. ERK7 orthologs have ciliogenesis-related roles in early branching unicellular lineages such as the parasitic apicomplexan *Toxoplasma gondii* as well as in the ciliated tissues of *Xenopus* embryos (Miyatake et al. 2015, O’Shaughnessy et al. 2020). The social amoeba *Dictyostelium discoideum* also has two MAPKs, one of which is orthologous to human ERK7 (Goldberg et al. 2006) and necessary for chemotaxis (Schwebs et al. 2018, Nichols et al. 2019). ERK7 orthologs were not identified among non-animal opisthokonts in this study. Interestingly, it has also been noted that ERK7 orthologs are missing in organisms that do not produce cilia or flagella during any point in development (O’Shaughnessy et al. 2020). Together this suggests that ERK7 is different from most of the other MAPKs that are responsible for fundamental cell physiology.

ERK7 has no known upstream MAP2K or MAP3K regulator (Cargnello and Roux, 2011) and is activated by autophosphorylation. Some ERK7 orthologs have a derived substitution of glutamate to aspartate (T-D-Y), the other amino acid with a negatively charged side chain. The sole exception to a negatively charged central residue in ERK7 is the valine seen in apusomonadid *Thecamonas trahens*. The functional relevance of this substitution is unknown. The conservation of the original T-E/D-Y phosphorylation motif nevertheless suggests that it is essential for ERK7 function.

The atypical human ERK3/4 has an unconventional S-E-G motif (Figure 6B). Through ortholog identification in diverging taxa, we were able to trace the origin of this unusual motif variation. We find that in sponge *Amphimedon queenslandica*, one additional residue was inserted into this motif, resulting in a T-D-S-Y variant. In other species representing early animal divergences, the first phosphorylation site is conserved but the following residues are highly variable, implying loss of the second site. Together, this represents the single occasion across the MAPK radiation where this highly conserved duo phosphorylation motif disintegrated. On the stem of Chordata (540-550 mya), the remaining threonine was replaced by serine, a phosphoacceptor less preferred by Ser/Thr kinases. On the stem of Vertebrata, the S-E-G motif achieves fixation. The trend towards decreased attractiveness to MAP2Ks may reflect the gradual uncoupling of ERK3/4 from upstream activation.

In our reconstructed phylogeny, the divergence of *_MRCA_*ERK1/2 and *_MRCA_*ERK3/4 occurs no later than the stem of Choanozoa. Choanoflagellates have one ortholog each of ERK1/2 and ERK3/4, both with the primitive T-E-Y motif. The fixation of the S-E-G phosphorylation motif however suggests that functional changes in *_MRCA_*ERK3/4 occurred on the stem vertebrate lineage. If so, the emergence of *_MRCA_*ERK3/4 significantly predates the derived atypical function of ERK3/4. This raises the possibility that *_MRCA_*ERK3/4 may be functionally redundant to *_MRCA_*ERK1/2 or nonfunctional, and thus easier lost due to their non-vital role. Indeed, multiple non-vertebrate species appear to have lost ERK3/4 orthologs.

The substitutions traced in the ERK3/4 motif may also provide insight into the phylogeny of NLK, which we excluded from our tree due to the phylogenetic implausibility of its initial reconstructed position, basal to all the other non-ERK7 homologs. NLK orthologs were identified across Animalia except for Ctenophora, but we did not find any non-animal orthologs. Analysis of the NLK phosphorylation motif revealed the fixation of a T-Q-E variant prior to the divergence of Cnidaria and Bilateria. This change was preceded by a transitional T-x-x motif that lost the second phosphorylation site. Using parsimony as the baseline arbiter of primary homology (Hennig’s Auxiliary Principle; see Hennig 1966, Thiele 1993), NLK would be considered the sister paralog of ERK3/4, the only other MAPK paralog that transitioned through a T-x-x motif. However, it is also probable that loss of the second phosphoacceptor in ERK3/4 and NLK are independent events.

Previously, the atypical MAPKs were proposed to be basal divergences that preserved an autoregulated primordial state (Sang et al. 2019). The relatively late emergence of *_MRCA_*ERK3/4 and NLK however suggests that the atypical MAPKs are a polyphyletic group. Retracing alterations in the dual phosphorylation motif further lends credence to the hypothesis that the autoregulation of some atypical MAPKs is a homoplastic feature that may have evolved more than once. In particular, the conserved ancestral motif in ERK7 and the deviant motifs of ERK3/4 and NLK suggest that atypical MAPKs attained independence from MAP2K activation by different mechanisms. Overall, our inference is that *_MRCA_*MAPK was not autoregulated.

### Links to the evolution of animal multicellularity

Human TKL MAP3Ks are often associated with cell-cell signaling molecules and growth factors downstream of GPCRs or receptor tyrosine kinases, which expanded dramatically in the path to multicellular animals (Suga et al. 2012, de Mendoza et al. 2014, Peterson et al. 2022). The downstream functions of TKLs are also tightly associated with processes key to organismal tissue control, including apoptosis and cell proliferation. In particular, the two TKL expansion events on the stem of Animalia are associated with functions critical to animal multicellularity. First, the MLK-JNK expansion produced a pathway dedicated to regulating programmed cell death. Second, the divergence of KSRs from the RAFs, which are necessary for enhancing RAF activity and (Clapéron and Therrien 2007), have critical regulatory roles for cell proliferation and fate regulation in both *Drosophila* and *Caenorhabditis* embryogenesis (Therrien et al. 1996, Rocheleau et al. 2008). Overall, the expansion of TKL MAPKs at the base of the animal radiation may have played a crucial role in facilitating animal multicellularity. Many studies emphasize the acquisition of functionally novel ‘key innovations’ to identify mechanisms underlying evolutionary success (Miller et al. 2023). The dramatic TKL MAP3K expansion at the base of Animalia, however, reflects a regulatory refinement that permitted the boosting and finetuning of deeply primitive cellular processes.

### Limitations and Future Direction

In this study, we reconstruct the evolutionary history of the human MAPK signaling network through meticulous identification of orthologs along the human phylogenetic backbone. Some questions cannot be confidently answered with our approach. Any divergence events outside the human evolutionary backbone would not be captured by our sampling strategy, thus limiting our ability to establish orthology among non-human homologs. We also have phylogenetic conflicts that we were unable to reconcile, especially as the conserved kinase domain contains only hundreds of loci. Tree conflicts are difficult to wholly eliminate, even at resolutions provided by transcriptome-based phylogenetic analysis (Sharma et al. 2014). In particular, the precision and resolution of our phylogenic reconstruction is limited by the paucity of high-quality annotated transcriptomes currently available, especially for non-model organisms that occupy critical phylogenetic positions. For these reasons, we refrain from absolute numerical dating of divergence events. Such estimates can be more misleading than informative in an incomplete paralog subtree where one branch can represent more than a billion years of evolution.

Phylogenetic relationships between homologs can be inferred from sequence alignments. However, experimental phenotypic assessment of homolog behavior in distant and closely related species is critical for determining polarity (i.e., plesiomorphic versus apomorphic) of associated functional traits and for phylogenetic bracketing of ancestral phenotypes. That the majority of molecular and genetic functional studies are conducted in a small number of established model organisms makes this challenging. In our study, the sequenced genomes of basal lineages Breviatea and Apusomonadida were critical for establishing the initial framework of the MAPK signaling network, but functional insights could only be gleaned from a few rare studies. This underscores the need for more functional studies and genome sequencing in non-standard organisms.

## Conclusion

A comprehensive understanding of the MAPK signaling network requires deciphering its deep evolutionary history. Our phylogenetic analyses of all three levels of this signaling network recovered patterns from which major evolutionary events could be inferred. With this inclusive approach, we characterize the MRCAs of all MAPK signaling pathways currently established in humans. Our synthesis of inferred phylogenetic patterns with published experimental data across diverse eukaryotic systems allowed the generation of high-resolution phylogenetic models for the evolution of molecular networks. Our results show distinct differences in roles of STE and TKL MAP3K in the elaboration of MAPK signaling: STE MAP3Ks are ancestral pathway components that maintained their essential functions throughout eukaryotic history, whereas TKL MAP3Ks were integrated later but were key drivers of network expansion. We propose that the complexity of the modern MAPK signaling network was established via functional specialization and increased specificity rather than by the acquisition of innovative functions. Our framework for signaling network evolution provides ground for evolutionary and functional hypotheses regarding the origin of novel pathways and their subsequent refinement and maintenance.

## Acknowledgements.

We thank Don Cerio, Jacob Wilson, and Will Foster for providing comments during various discussions. We also thank JE Lin for providing the illustrations of various eukaryotes. We acknowledge our funding sources: NIGMS R35GM133499, NCI R01CA279546, and NSF CAREER 1844994 to SR.

## Author contributions

Conceptualization: EJ Huang (EJH), Amy F. Peterson (AFP), Sergi Regot (SR), Gabriel S. Bever (GSB); Data Curation: EJH, Fernando Torres (FT); Formal Analysis: EJH, FT; Investigation: EJH, Jeeun Parksong (JP), FT; Methodology: EJH; Project Administration: EJH, SR, GSB; Resources: EJH, SR, GSB; Software: EJH, FT; Supervision: SR, GSB; Validation: EJH, JP, AFP, FT; Visualization: EJH; Writing – Original Draft Preparation: EJH, JP, GSB; Writing – Review & Editing: EJH, JP, AFP, FT, SR, GSB

## Declaration of Interests

The authors declare no competing interests.

## Notes

### Competing Interest Statement

The authors have declared no competing interest.

## References

Asai, T., Tena, G., Plotnikova, J., Willmann, M. R., Chiu, W., Gomez-Gomez, L., Boller, T., Ausubel, F. M., & Sheen, J. (2002). MAP kinase signalling cascade in *Arabidopsis* innate immunity. Nature (London), 415(6875), 977–983. 10.1038/415977a

Bardwell, L. (2006). Mechanisms of MAPK signalling specificity. Biochemical Society Transactions, 34, 837–841. 10.1042/BST0340837

Bergsten, J. (2005). A review of long-branch attraction. Cladistics, 21(2), 163–193. 10.1111/j.1096-0031.2005.00059.x

Betts, H. C., Puttick, M. N., Clark, J. W., Williams, T. A., Donoghue, P. C. J., & Pisani, D. (2018). Integrated genomic and fossil evidence illuminates life’s early evolution and eukaryote origin. Nature Ecology & Evolution, 2(10), 1556–1562. 10.1038/s41559-018-0644

Böhm, M., Gamulin, V., Schröder, H. C., & Müller, W. E. G. (2002). Evolution of osmosensing signal transduction in Metazoa: Stress-activated protein kinases p38 and JNK. Cell and Tissue Research, 308(3), 431–438. 10.1007/s00441-002-0535-x

Bridge, D., Theofiles, A. G., Holler, R. L., Marcinkevicius, E., Steele, R. E., & Martínez, D. E. (2010). FoxO and stress responses in the cnidarian *Hydra vulgaris*. PLoS ONE, 5(7), e11686. 10.1371/journal.pone.0011686

Brown, M. W., Sharpe, S. C., Silberman, J. D., Heiss, A. A., Lang, B. F., Simpson, A. G. B., & Roger, J. (2013). Phylogenomics demonstrates that breviate flagellates are related to opisthokonts and apusomonads. *Proceedings of the Royal Society B*, Biological Sciences, 280(1769), 20131755. 10.1098/rspb.2013.1755

Buchmann, K. (2014). Evolution of innate immunity: Clues from invertebrates via fish to mammals. Frontiers in Immunology, 5, 459. 10.3389/fimmu.2014.00459

Caffrey, D. R., O’Neill, L. A., & Shields, D. C. (1999). The evolution of the MAP kinase pathways: Coduplication of interacting proteins leads to new signaling cascades. Journal of Molecular Evolution, 49(5), 567–582. 10.1007/PL00006578

Canagarajah, B. J., Khokhlatchev, A., Cobb, M. H., & Goldsmith, E. J. (1997). Activation mechanism of the MAP kinase ERK2 by dual phosphorylation. Cell, 90(5), 859–869. 10.1016/S0092-8674(00)80351-7

Cantino, P. D., & Queiroz, K. d. (2020). PhyloCode: A phylogenetic code of biological nomenclature. CRC Press.

Capella-Gutierrez, S., Silla-Martinez, J., & Gabaldon, T. (2009). trimAl: A tool for automated alignment trimming in large-scale phylogenetic analyses. Bioinformatics, 25(15), 1972–1973. 10.1093/bioinformatics/btp348

Cargnello, M., & Roux, P. P. (2011). Activation and function of the MAPKs and their substrates, the MAPK-activated protein kinases. Microbiology and Molecular Biology Reviews, 75(1), 50–83. 10.1128/MMBR.00031-10

Clapéron, A., & Therrien, M. (2007). KSR and CNK: Two scaffolds regulating RAS-mediated RAF activation. Oncogene, 26(22), 3143–3158. 10.1038/sj.onc.1210408

Clark, K., Karsch-Mizrachi, I., Lipman, D. J., Ostell, J., & Sayers, E. W. (2016). GenBank. Nucleic Acids Research, 44, D67–D72. 10.1093/nar/gkv1276

de Mendoza, A., Sebé-Pedrós, A., & Ruiz-Trillo, I. (2014). The evolution of the GPCR signaling system in eukaryotes: Modularity, conservation, and the transition to metazoan multicellularity. Genome Biology and Evolution, 6(3), 606–619. 10.1093/gbe/evu038

de Mendoza, A., Suga, H., Permanyer, J., Irimia, M., & Ruiz-Trillo, I. (2015). Complex transcriptional regulation and independent evolution of fungal-like traits in a relative of animals. eLife, 4, e08904. 10.7554/eLife.08904

de Nadal, E., & Posas, F. (2022). The HOG pathway and the regulation of osmoadaptive responses in yeast. FEMS Yeast Research, 22(1) 10.1093/femsyr/foac013

Dehal, P., & Boore, J. L. (2005). Two rounds of whole genome duplication in the ancestral vertebrate. PLoS Biology, 3(10), e314. 10.1371/journal.pbio.0030314

Déléris, P., Trost, M., Topisirovic, I., Tanguay, P., Borden, K. L. B., Thibault, P., & Meloche, S. (2011). Activation loop phosphorylation of ERK3/ERK4 by group I p21-activated kinases (PAKs) defines a novel PAK-ERK3/4-MAPK-activated protein kinase 5 signaling pathway. Journal of Biological Chemistry, 286(8), 6470–6478. 10.1074/jbc.M110.181529

Deniz, O., Hasygar, K., & Hietakangas, V. (2023). Cellular and physiological roles of the conserved atypical MAP kinase ERK7. FEBS Letters, 597(5), 601–607. 10.1002/1873-3468.14521

Dóczi, R., Ökrész, L., Romero, A. E., Paccanaro, A., & Bögre, L. (2012). Exploring the evolutionary path of plant MAPK networks. Trends in Plant Science, 17(9), 518–525. 10.1016/j.tplants.2012.05.009

Drew, B. A., Burow, M. E., & Beckman, B. S. (2012). MEK5/ERK5 pathway: The first fifteen years. Biochimica et Biophysica Acta, 1825(1), 37–48. 10.1016/j.bbcan.2011.10.002

Duch, A., de Nadal, E., & Posas, F. (2012). The p38 and Hog1 SAPKs control cell cycle progression in response to environmental stresses. FEBS Letters, 586(18), 2925–2931. 10.1016/j.febslet.2012.07.034

Dudin, O., Ondracka, A., Grau-Bové, X., Haraldsen, A. A., Toyoda, A., Suga, H., Bråte, J., & Ruiz-Trillo, I. Transcriptome - Sphaeroforma arctica. 10.6084/m9.figshare.8299529.v2

Eme, L., Tamarit, D., Caceres, E. F., Stairs, C. W., De Anda, V., Schön, M. E., Seitz, K. W., Dombrowski, N., Lewis, W. H., Homa, F., Saw, J. H., Lombard, J., Nunoura, T., Li, W., Hua, Z., Chen, L., Banfield, J. F., John, E. S., Reysenbach, A., . . . Ettema, T. J. G. (2023). Inference and reconstruction of the heimdallarchaeial ancestry of eukaryotes. Nature (London), 618(7967), 992–999. 10.1038/s41586-023-06186-2

Flajnik, M. F., & Kasahara, M. (2010). Origin and evolution of the adaptive immune system: Genetic events and selective pressures. Nature Reviews Genetics, 11(1), 47–59. 10.1038/nrg2703

Fraser, H. B., Hirsh, A. E., Wall, D. P., Eisen, M. B., & Li, W. (2004). Coevolution of gene expression among interacting proteins. Proceedings of the National Academy of Sciences, 101(24), 9033–9038. 10.1073/pnas.0402591101

Gantke, T., Sriskantharajah, S., & Ley, S. C. (2011). Regulation and function of TPL-2, an IκB kinase-regulated MAP kinase kinase kinase. Cell Research, 21(1), 131–145. 10.1038/cr.2010.173

Goldberg, J. M., Manning, G., Liu, A., Fey, P., Pilcher, K. E., Xu, Y., & Smith, J. L. (2006). The dictyostelium Kinome—Analysis of the protein kinases from a simple model organism. PLoS Genetics, 2(3), e38. 10.1371/journal.pgen.0020038

González-Rubio, G., Sellers-Moya, Á, Martín, H., & Molina, M. (2021). A walk-through MAPK structure and functionality with the 30-year-old yeast MAPK Slt2. International Microbiology, 24(4), 531–543. 10.1007/s10123-021-00183-z

Guo, X., Wang, Z., Liu, L., & Li, Y. (2021). Transcriptome and metabolome analyses of cold and darkness-induced pellicle cysts of *Scrippsiella trochoidea*. BMC Genomics, 22(1), 526. 10.1186/s12864-021-07840-7

Hadwiger, J. A., & Nguyen, H. (2011). MAPKs in development: Insights from *Dictyostelium* signaling pathways. Biomolecular Concepts, 2(1-2), 39–46. 10.1515/bmc.2011.004

Hennig, W. (1966). Phylogenetic systematics. Univ. of Illinois Press.

Herskowitz, I. (1995). MAP kinase pathways in yeast: For mating and more. Elsevier Inc. 10.1016/0092-8674(95)90402-6

Horton, A. A., Wang, B., Camp, L., Price, M. S., Arshi, A., Nagy, M., Nadler, S. A., Faeder, J. R., & Luckhart, S. (2011). The mitogen-activated protein kinome from *Anopheles gambiae*: Identification, phylogeny and functional characterization of the ERK, JNK and p38 MAP kinases. BMC Genomics, 12(1), 574. 10.1186/1471-2164-12-574

Ikeda, A., Masaki, M., Kozutsumi, Y., Oka, S., & Kawasaki, T. (2001). Identification and characterization of functional domains in a mixed lineage kinase LZK. FEBS Letters, 488(3), 190–195. 10.1016/S0014-5793(00)02432-7

Jin, H., Axtell, M. J., Dahlbeck, D., Ekwenna, O., Zhang, S., Staskawicz, B., & Baker, B. (2002). NPK1, an MEKK1-like mitogen-activated protein kinase kinase kinase, regulates innate immunity and development in plants. Developmental Cell, 3(2), 291–297. 10.1016/S1534-5807(02)00205-8

Jo, M., Chung, A. Y., Yachie, N., Seo, M., Jeon, H., Nam, Y., Seo, Y., Kim, E., Zhong, Q., Vidal, M., Park, H. C., Roth, F. P., & Suk, K. (2017). Yeast genetic interaction screen of human genes associated with amyotrophic lateral sclerosis: Identification of MAP2K5 kinase as a potential drug target. Genome Research, 27(9), 1487–1500. 10.1101/gr.211649.116

Johnson, M., Zaretskaya, I., Raytselis, Y., Merezhuk, Y., McGinnis, S., & Madden, T. L. (2008). NCBI BLAST: A better web interface. Nucleic Acids Research, 36(-2), W5-W9. 10.1093/nar/gkn201

Katoh, K., & Standley, D. M. (2013). MAFFT multiple sequence alignment software version 7: Improvements in performance and usability. Molecular Biology and Evolution, 30(4), 772–780. 10.1093/molbev/mst010

Kenny, N. J., Francis, W. R., Rivera-Vicéns, R. E., Juravel, K., de Mendoza, A., Díez-Vives, C., Lister, R., Bezares-Calderón, L. A., Grombacher, L., Roller, M., Barlow, L. D., Camilli, S., Ryan, J. F., Wörheide, G., Hill, A. L., Riesgo, A., & Leys, S. P. (2020). Tracing animal genomic evolution with the chromosomal-level assembly of the freshwater sponge *Ephydatia muelleri*. Nature Communications, 11(1), 3676. 10.1038/s41467-020-17397-w

Krieger, F., Möglich, A., & Kiefhaber, T. (2005). Effect of proline and glycine residues on dynamics and barriers of loop formation in polypeptide chains. Journal of the American Chemical Society, 127(10), 3346–3352. 10.1021/ja042798i

Kultz, D. (., B. (1998). Phylogenetic and functional classification of mitogen- and stress-activated protein kinases. Journal of Molecular Evolution, 46(5), 571–588. 10.1007/PL00006338

Kyriakis, J. M., & Avruch, J. (2012). Mammalian MAPK signal transduction pathways activated by stress and inflammation: A 10-year update. Physiological Reviews, 92(2), 689–737. 10.1152/physrev.00028.2011

Multicellgenome Lab, & Torruella, G. (2017a). Transcriptome - Nutomonas longa. 10.6084/m9.figshare.4560862.v2

Multicellgenome Lab, & Torruella, G. (2017b). Transcriptome - Parvularia atlantis. 10.6084/m9.figshare.3898485.v4

Murakawa, Y., Valter, S., Barr, H., London, N., & Kinoshita, T. (2020). Structural basis for producing selective MAP2K7 inhibitors. Bioorganic & Medicinal Chemistry Letters, 30(22), 127546. 10.1016/j.bmcl.2020.127546

Nakatani, Y., Shingate, P., Ravi, V., Pillai, N. E., Prasad, A., McLysaght, A., & Venkatesh, B. (2021). Reconstruction of proto-vertebrate, proto-cyclostome and proto-gnathostome genomes provides new insights into early vertebrate evolution. Nature Communications, 12(1), 4489. 10.1038/s41467-021-24573-z

Lal, H., Ahmad, F., Parikh, S., & Force, T. (2014). Troponin I-interacting protein kinase: A novel cardiac-specific kinase, emerging as a molecular target for the treatment of cardiac disease. Circulation Journal, 78(7), 1514–1519. 10.1253/circj.CJ-14-0543

Lewis, T. S., Shapiro, P. S., & Ahn, N. G. (1998). Signal transduction through MAP kinase cascades. Advances in cancer research (pp. 49–139). Elsevier Science & Technology. 10.1016/s0065-230x(08)60765-4

Li, M., Liu, J., & Zhang, C. (2011). Evolutionary history of the vertebrate mitogen activated protein kinases family. PLoS ONE, 6(10), e26999. 10.1371/journal.pone.0026999

Maeda, T., Wurgler-Murphy, S., & Saito, H. (1994). A two-component system that regulates an osmosensing MAP kinase cascade in yeast. Nature (London), 369(6477), 242. https://www.ncbi.nlm.nih.gov/pubmed/8183345

Manning, G., Whyte, D. B., Martinez, R., Hunter, T., & Sudarsanam, S. (2002). The protein kinase complement of the human genome. Science, 298(5600), 1912–1934. 10.1126/science.1075762

Miller, A. H., Stroud, J. T., & Losos, J. B. (2023). The ecology and evolution of key innovations. Trends in Ecology & Evolution (Amsterdam), 38(2), 122–131. 10.1016/j.tree.2022.09.005

Miyatake, K., Kusakabe, M., Takahashi, C., & Nishida, E. (2015). ERK7 regulates ciliogenesis by phosphorylating the actin regulator CapZIP in cooperation with Dishevelled. Nature Communications, 6(1), 6666. 10.1038/ncomms7666

Moreland, R. T., Nguyen, A., Ryan, J. F., & Baxevanis, A. D. (2020). The *Mnemiopsis* genome project portal: Integrating new gene expression resources and improving data visualization. Database: The Journal of Biological Databases and Curation, 2020. 10.1093/database/baaa029

Nguyen, A., Burack, W. R., Stock, J. L., Kortum, R., Chaika, O. V., Afkarian, M., Muller, W. J., Murphy, K. M., Morrison, D. K., Lewis, R. E., McNeish, J., & Shaw, A. S. (2002). Kinase suppressor of Ras (KSR) is a scaffold which facilitates mitogen-activated protein kinase activation in vivo. Molecular and Cellular Biology, 22(9), 3035–3045. 10.1128/MCB.22.9.3035-3045.2002

Nichols, J. M. E., Paschke, P., Peak-Chew, S., Williams, T. D., Tweedy, L., Skehel, M., Stephens, E., Chubb, J. R., & Kay, R. R. (2019). The atypical MAP kinase ErkB transmits distinct chemotactic signals through a core signaling module. Developmental Cell, 48(4), 491–505. 10.1016/j.devcel.2018.12.001

Nithianandarajah-Jones, G., Wilm, B., Goldring, C. E. P., Müller, J., & Cross, M. J. (2012). ERK5: Structure, regulation and function. Cellular Signalling, 24(11), 2187–2196. 10.1016/j.cellsig.2012.07.007

O’Shaughnessy, W. J., Hu, X., Beraki, T., McDougal, M., & Reese, M. L. (2020). Loss of a conserved MAPK causes catastrophic failure in assembly of a specialized cilium-like structure in toxoplasma gondii. Molecular Biology of the Cell, 31(9), 881–888. 10.1091/mbc.E19-11-0607

Pang, K., Wang, W., Qin, J., Shi, Z., Hao, L., Ma, Y., Xu, H., Wu, Z., Pan, D., Chen, Z., & Han, C. (2022). Role of protein phosphorylation in cell signaling, disease, and the intervention therapy. MedComm (2020), *3*(4), e175. 10.1002/mco2.175

Paps, J., & Holland, P. W. H. (2018). Reconstruction of the ancestral metazoan genome reveals an increase in genomic novelty. Nature Communications, 9(1), 1730–1738. 10.1038/s41467-018-04136-5

Park, H. L., Seo, D. H., Lee, H. Y., Bakshi, A., Park, C., Chien, Y., Kieber, J. J., Binder, B. M., & Yoon, G. M. (2023). Ethylene-triggered subcellular trafficking of CTR1 enhances the response to ethylene gas. Nature Communications, 14(1), 365. 10.1038/s41467-023-35975-6

Paudel, R., Fusi, L., & Schmidt, M. (2021). The MEK5/ERK5 pathway in health and disease. International Journal of Molecular Sciences, 22(14), 7594. 10.3390/ijms22147594

Pereira, D. M., & Rodrigues, C. M. P. (2020). Targeted avenues for cancer treatment: The MEK5– ERK5 signaling pathway. Trends in Molecular Medicine, 26(4), 394–407. 10.1016/j.molmed.2020.01.006

Peterson, A. F., Ingram, K., Huang, E. J., Parksong, J., McKenney, C., Bever, G. S., & Regot, S. (2022). Systematic analysis of the MAPK signaling network reveals MAP3K-driven control of cell fate. Cell Systems, 13(11), 885–894.e4. 10.1016/j.cels.2022.10.003

Roberts, R. L., & Fink, G. R. (1994). Elements of a single MAP kinase cascade in saccharomyces cerevisiae mediate two developmental programs in the same cell type: Mating and invasive growth. Genes & Development, 8(24), 2974–2985. 10.1101/gad.8.24.2974

Robitaille, H., Simard-Bisson, C., Larouche, D., Tanguay, R. M., Blouin, R., & Germain, L. (2010). The small heat-shock protein Hsp27 undergoes ERK-dependent phosphorylation and redistribution to the cytoskeleton in response to dual leucine zipper-bearing kinase expression. Journal of Investigative Dermatology, 130(1), 74–85. 10.1038/jid.2009.185

Rocheleau, C. E., Cullison, K., Huang, K., Bernstein, Y., Spilker, A. C., & Sundaram, M. V. (2008). The *Caenorhabditis elegans ekl* (enhancer of ksr-1 lethality) genes include putative components of a germline small RNA pathway. Genetics, 178(3), 1431–1443. 10.1534/genetics.107.084608

Rosindell, J., & Harmon, L. J. (2012). OneZoom: A fractal explorer for the tree of life. PLoS Biology, 10(10), e1001406. 10.1371/journal.pbio.1001406

Samatar, A. A., & Poulikakos, P. I. (2014). Targeting RAS–ERK signalling in cancer: Promises and challenges. Nature Reviews Drug Discovery, 13(12), 928–942. 10.1038/nrd4281

Sang, D., Pinglay, S., Wiewiora, R. P., Selvan, M. E., Lou, H. J., Chodera, J. D., Turk, B. E., Gümüs, Z. H., & Holt, L. J. (2019). Ancestral reconstruction reveals mechanisms of ERK regulatory evolution. eLife, 8 10.7554/eLife.38805

Schwebs, D. J., Pan, M., Adhikari, N., Kuburich, N. A., Jin, T., & Hadwiger, J. A. (2018). *Dictyostelium* Erk2 is an atypical MAPK required for chemotaxis. Cellular Signalling, 46, 154–165. 10.1016/j.cellsig.2018.03.006

Shabardina, V., Charria Romero, P., Bercedo Saborido, G., Diaz-Mora, E., Cuenda, A., Ruiz Trillo, I., & Sanz-Ezquerro, J. (2023). Evolutionary analysis of p38 stress-activated kinases in unicellular relatives of animals suggests an ancestral function in osmotic stress. 10.1098/rsob.220314

Sharma, P. P., Kaluziak, S. T., Pérez-Porro, A. R., González, V. L., Hormiga, G., Wheeler, W. C., & Giribet, G. (2014). Phylogenomic interrogation of Arachnida reveals systemic conflicts in phylogenetic signal. Molecular Biology and Evolution, 31(11), 2963–2984. 10.1093/molbev/msu235

Smith, T. G., Sweetman, D., Patterson, M., Keyse, S. M., & Münsterberg, A. (2005). Feedback interactions between MKP3 and ERK MAP kinase control *scleraxis* expression and the specification of rib progenitors in the developing chick somite. Development, 132(6), 1305–1314. 10.1242/dev.01699

Stamatakis, A. (2014). RAxML version 8: A tool for phylogenetic analysis and post-analysis of large phylogenies. Bioinformatics, 30(9), 1312–1313. 10.1093/bioinformatics/btu033

Strassert, J. F. H., Irisarri, I., Williams, T. A., & Burki, F. (2021). A molecular timescale for eukaryote evolution with implications for the origin of red algal-derived plastids. Nature Communications, 12(1), 1879. 10.1038/s41467-021-22044-z

Suga, H., Dacre, M., de Mendoza, A., Shalchian-Tabrizi, K., Manning, G., & Ruiz-Trillo, I. (2012). Genomic survey of premetazoans shows deep conservation of cytoplasmic tyrosine kinases and multiple radiations of receptor tyrosine kinases. Science Signaling, 5(222), ra35. 10.1126/scisignal.2002733

Swofford, D. L. (2003). PAUP^ phylogenetic analysis using parsimony (^and other methods). version 4. Sinauer Associates.

Takekawa, M., Posas, F., & Saito, H. (1997). A human homolog of the yeast Ssk2/Ssk22 MAP kinase kinase kinases, MTK1, mediates stress-induced activation of the p38 and JNK pathways. *EMBO Journal*, *16*(16), 4973-4982. 10.1093/emboj/16.16.4973

Therrien, M., Michaud, N. R., Rubin, G. M., & Morrison, D. K. (1996). KSR modulates signal propagation within the MAPK cascade. Genes & Development, 10(21), 2684–2695. 10.1101/gad.10.21.2684

Thiele, K. (1993). The holy grail of the perfect character: The cladistic treatment of morphometric data. Cladistics, 9(3), 275–304. 10.1006/clad.1993.1020

Thu, Y. M., & Richmond, A. (2010). NF-κB inducing kinase: A key regulator in the immune system and in cancer. Cytokine & Growth Factor Reviews, 21(4), 213–226. 10.1016/j.cytogfr.2010.06.002

Tusa, I., Gagliardi, S., Tubita, A., Pandolfi, S., Urso, C., Borgognoni, L., Wang, J., Deng, X., Gray, N. S., Stecca, B., & Rovida, E. (2018). ERK5 is activated by oncogenic BRAF and promotes melanoma growth. Oncogene, 37(19), 2601–2614. 10.1038/s41388-018-0164-9

Ubersax, J. A., & Ferrell Jr, J. E. (2007). Mechanisms of specificity in protein phosphorylation. Nature Reviews Molecular Cell Biology, 8(7), 530–541. 10.1038/nrm2203

Van de Peer, Y., Maere, S., & Meyer, A. (2009). The evolutionary significance of ancient genome duplications. Nature Reviews Genetics, 10(10), 725–732. 10.1038/nrg2600

Weisberg, E., Parent, A., Yang, P. L., Sattler, M., Liu, Q., Liu, Q., Wang, J., Meng, C., Buhrlage, S. J., Gray, N., & Griffin, J. D. (2020). Repurposing of kinase inhibitors for treatment of COVID-19. Pharmaceutical Research, 37(9), 167. 10.1007/s11095-020-02851-7

You, B., Lee, M., Tien, N., Lee, M., Hsieh, H., Tseng, L., Chung, Y., & Lee, H. (2013). A novel approach to enhancing ganoderic acid production by *Ganoderma lucidum* using apoptosis induction. PLoS ONE, 8(1), e53616. 10.1371/journal.pone.0053616

Zaremba-Niedzwiedzka, K., Caceres, E. F., Saw, J. H., Bäckström, D., Juzokaite, L., Vancaester, E., Seitz, K. W., Anantharaman, K., Starnawski, P., Kjeldsen, K. U., Stott, M. B., Nunoura, T., Banfield, J. F., Schramm, A., Baker, B. J., Spang, A., & Ettema, T. J. G. (2017). *Asgard archaea* illuminate the origin of eukaryotic cellular complexity. Nature, 541(7637), 353–358. 10.1038/nature21031

Zhang, M., & Zhang, S. (2022). Mitogen-activated protein kinase cascades in plant signaling. Journal of Integrative Plant Biology, 64(2), 301–341. 10.1111/jipb.13215

Zheng, C. F., & Guan, K. L. (1994). Activation of MEK family kinases requires phosphorylation of two conserved ser/thr residues. The EMBO Journal, 13(5), 1123–1131. 10.1002/j.1460-2075.1994.tb06361.x

Zulawski, M., Schulze, G., Braginets, R., Hartmann, S., & Schulze, W. X. (2014). The *Arabidopsis* kinome: Phylogeny and evolutionary insights into functional diversification. BMC Genomics, 15(1), 548. 10.1186/1471-2164-15-548

