## Supplementary material for "Reconstructing the deep phylogeny of the MAPK signaling network: functional specialization via multi-tier coevolutionary expansion": SI Appendix

**This PDF file includes:**

Figures S1 to S10
Tables S1 to S4
SI References

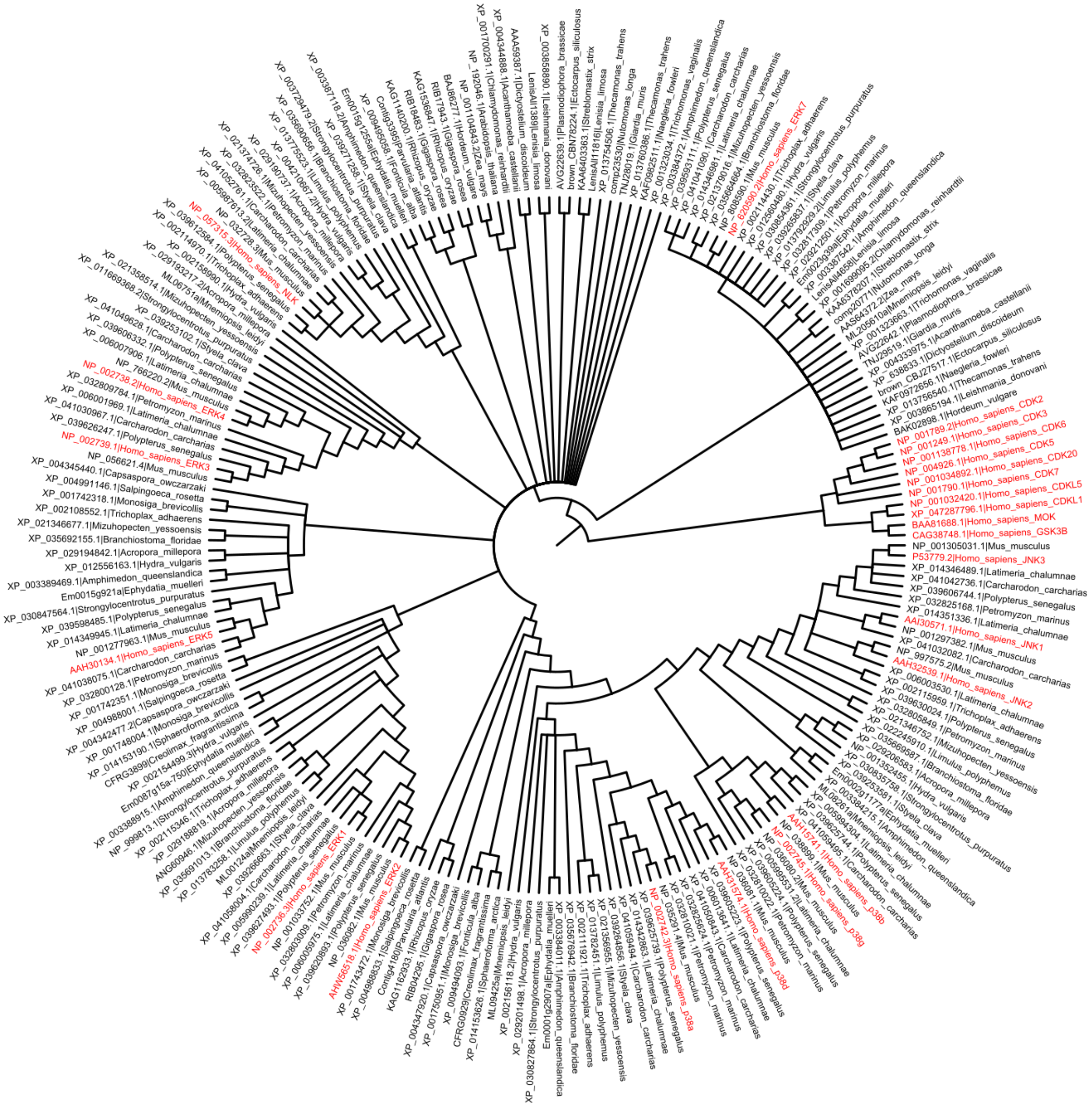

**Fig. S1. Strict consensus MAPK phylogeny.** Strict consensus MAPK cladogram of the ten maximum likelihood reconstruction replicates. The tree is rooted with human outgroup kinases. All human homologs are colored red and labeled with the names of corresponding proteins.

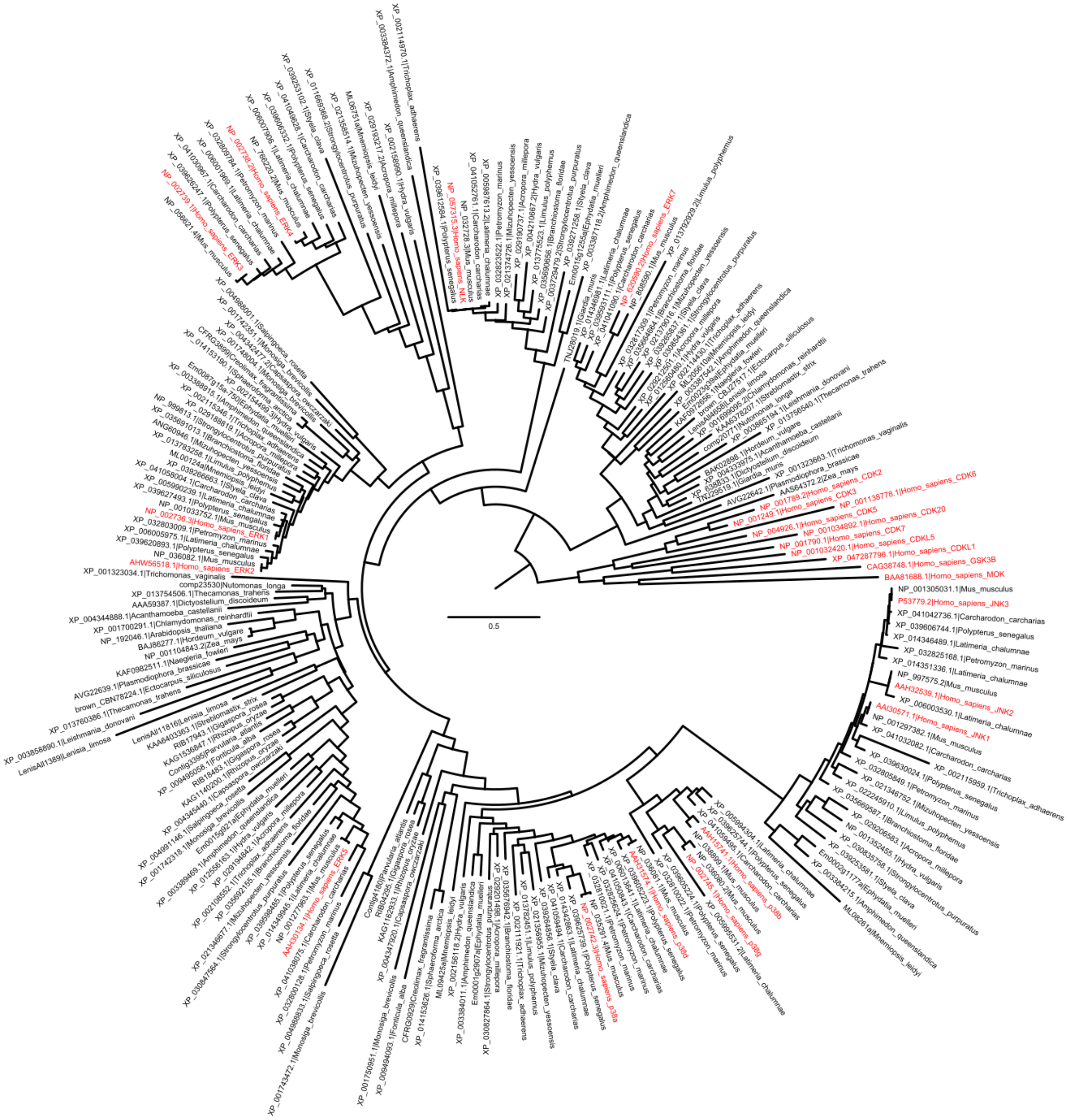

**Fig. S2. Maximum-likelihood MAPK phylogeny.** The MAPK phylogeny with the highest likelihood value among the ten technical replicates. The tree is rooted with human outgroup kinases. All human homologs are colored red and labeled with the names of corresponding proteins. Branch length represents the expected number of substitutions per unit time.

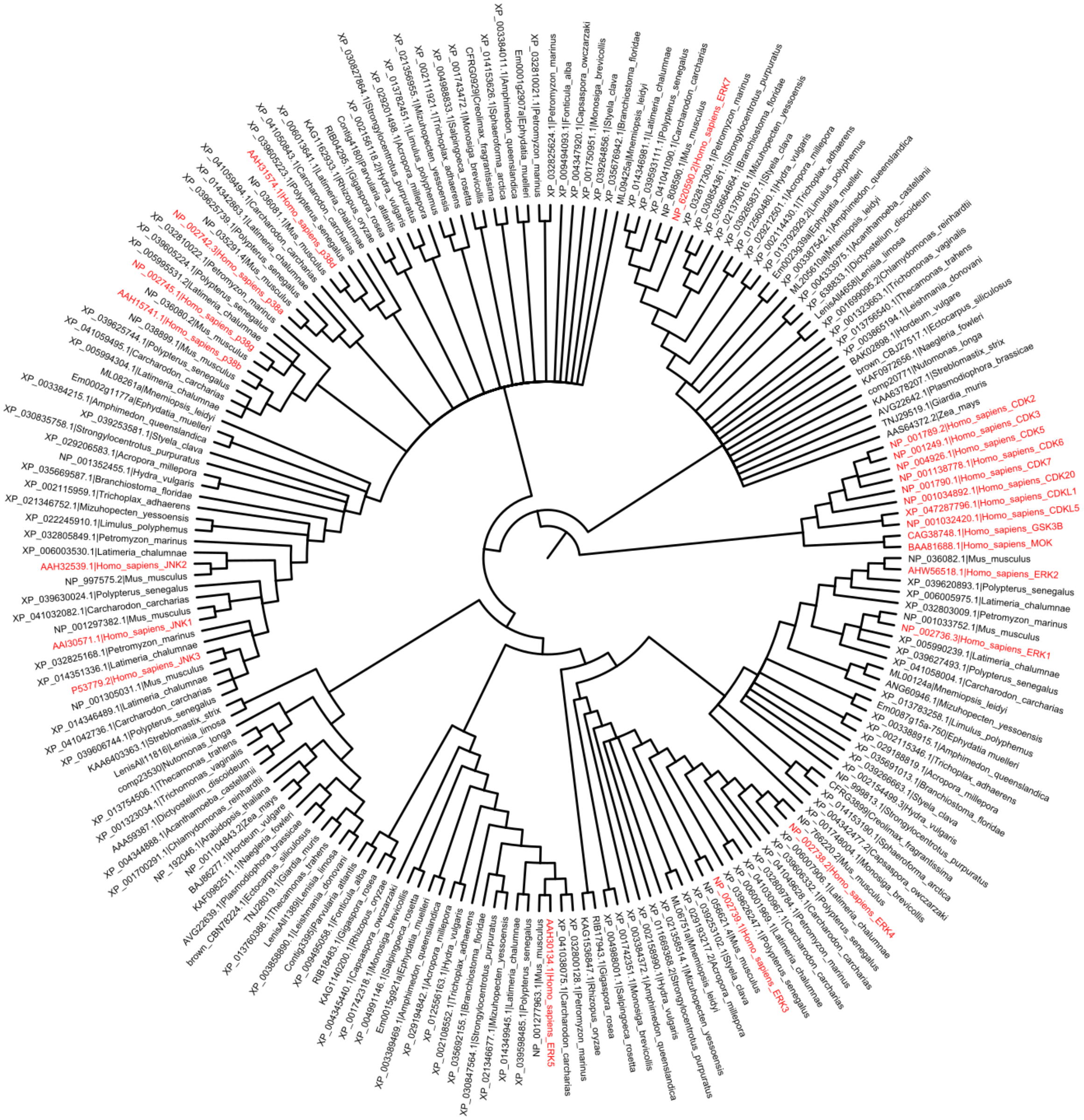

**Fig. S3. Strict consensus MAPK (without NLK) phylogeny.** Strict consensus MAPK cladogram (without NLK) of the ten maximum likelihood reconstruction replicates. The tree is rooted with human outgroup kinases. All human homologs are colored red and labeled with the names of corresponding proteins.

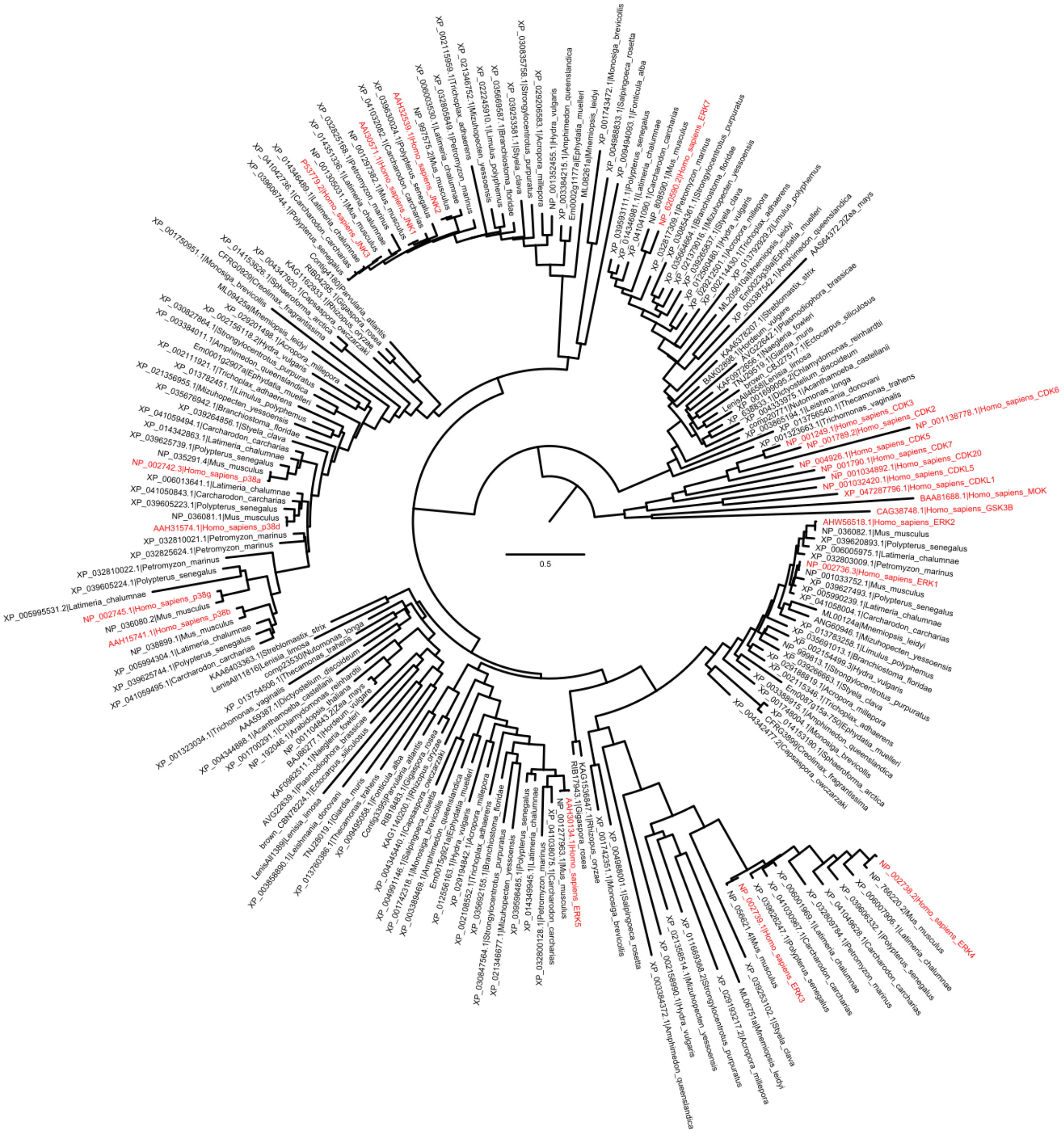


**Fig. S4. Maximum-likelihood MAPK (without NLK) phylogeny.** The MAPK (without NLK) phylogeny with the highest likelihood value among the ten technical replicates. The tree is rooted with human outgroup kinases. All human homologs are colored red and labeled with the names of corresponding proteins. Branch length represents the expected number of substitutions per unit time.

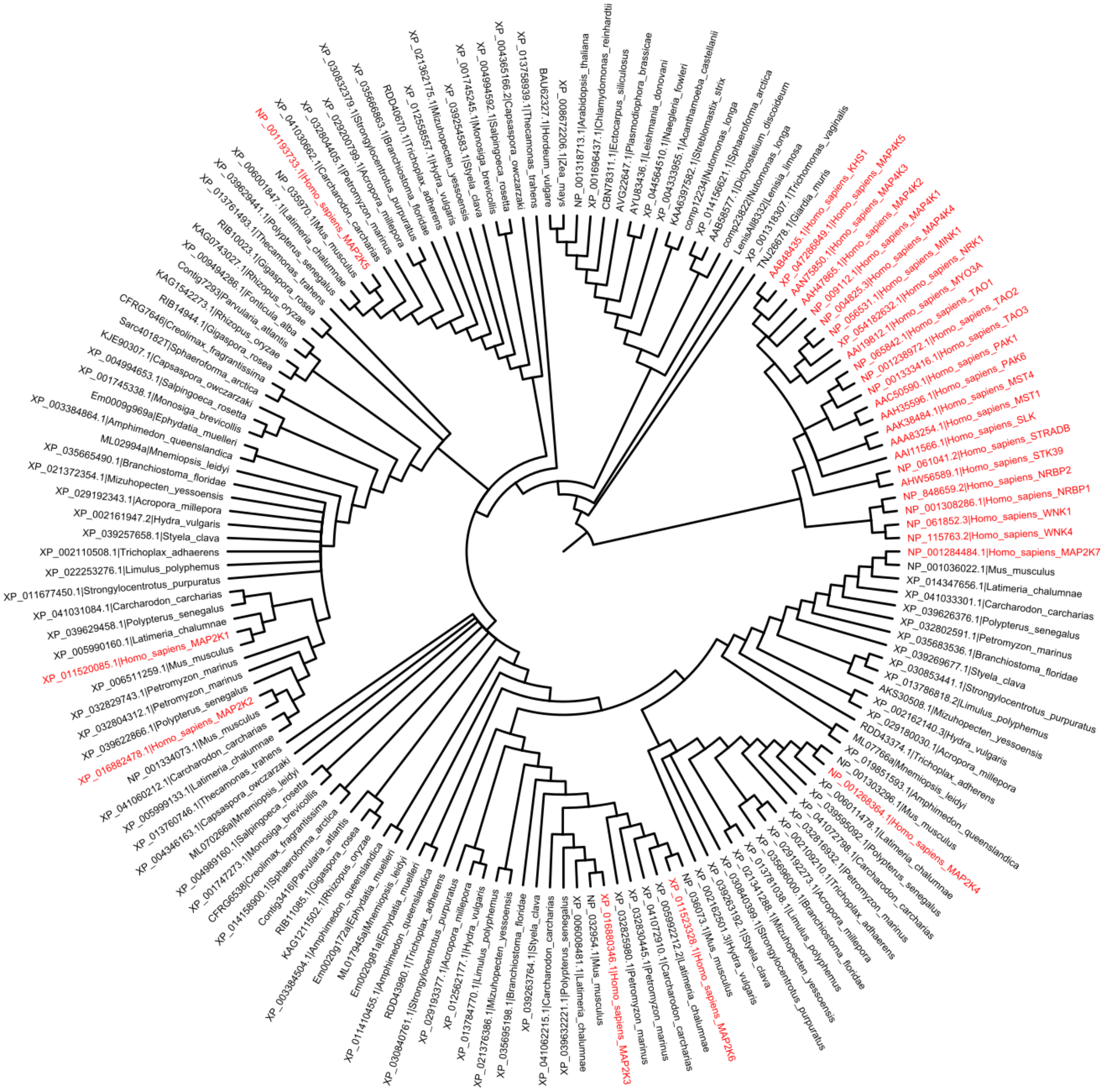

**Fig. S5. Strict consensus MAP2K phylogeny.** Strict consensus MAP2K cladogram of the ten maximum likelihood reconstruction replicates. The tree is rooted with human outgroup kinases. All human homologs are colored red and labeled with the names of corresponding proteins.

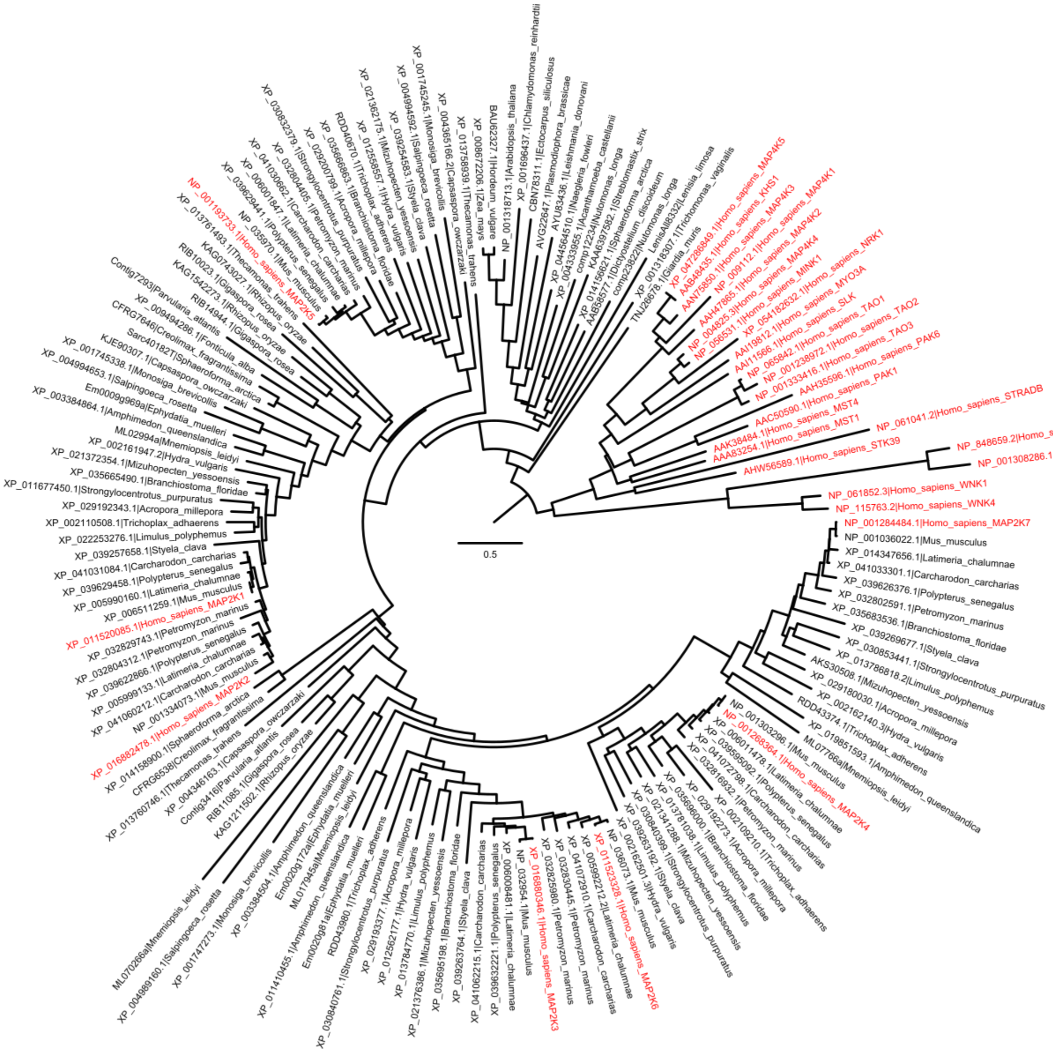


**Fig. S6. Maximum-likelihood MAP2K phylogeny.** The MAP2K phylogeny with the highest likelihood value among the ten technical replicates. The tree is rooted with human outgroup kinases. All human homologs are colored red and labeled with the names of corresponding proteins. Branch length represents the expected number of substitutions per unit time.

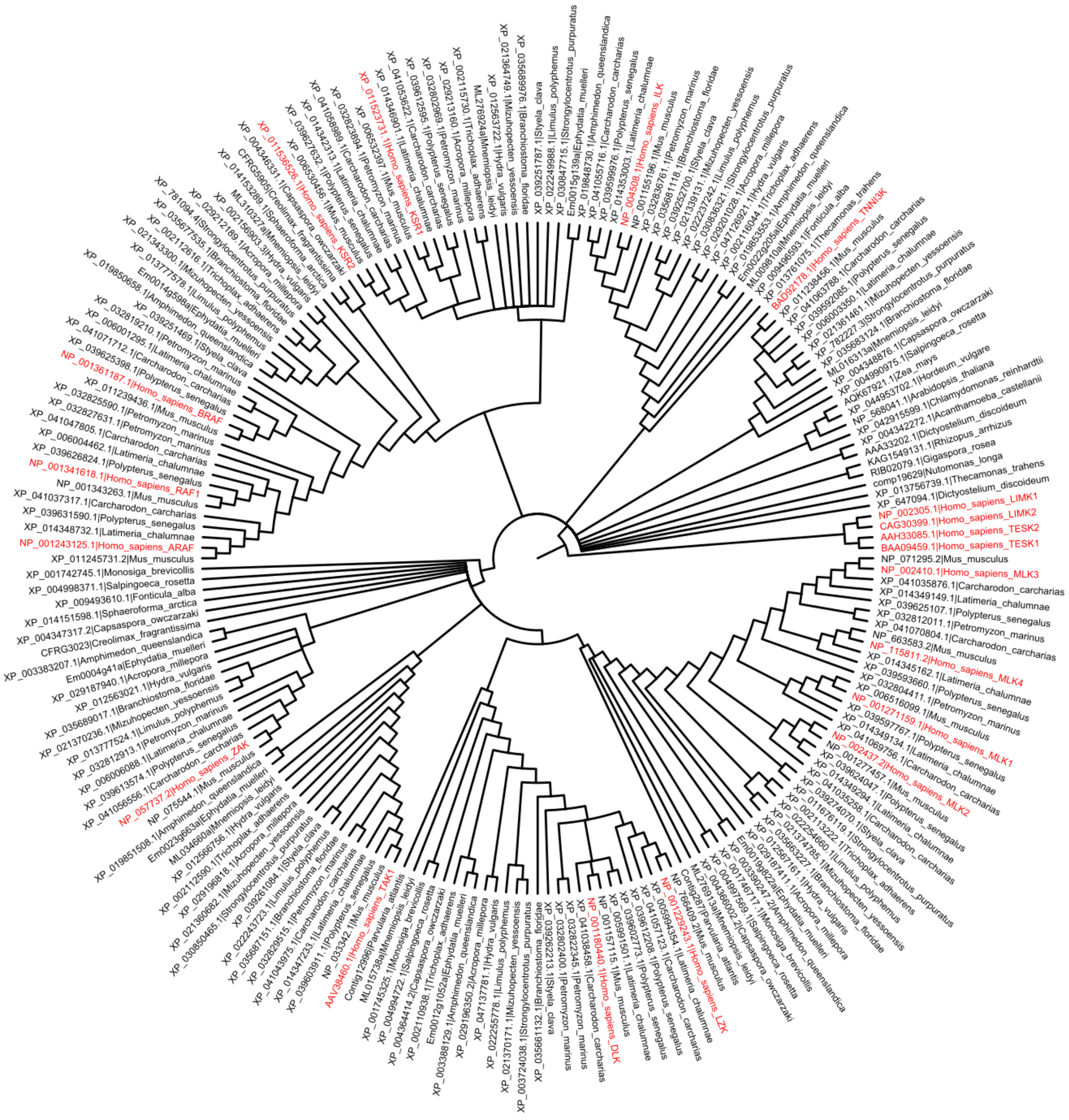

**Fig. S7. Strict consensus TKL MAP3K phylogeny.** Strict consensus TKL MAP3K cladogram of the ten maximum likelihood reconstruction replicates. The tree is rooted with human outgroup kinases. All human homologs are colored red and labeled with the names of corresponding proteins.

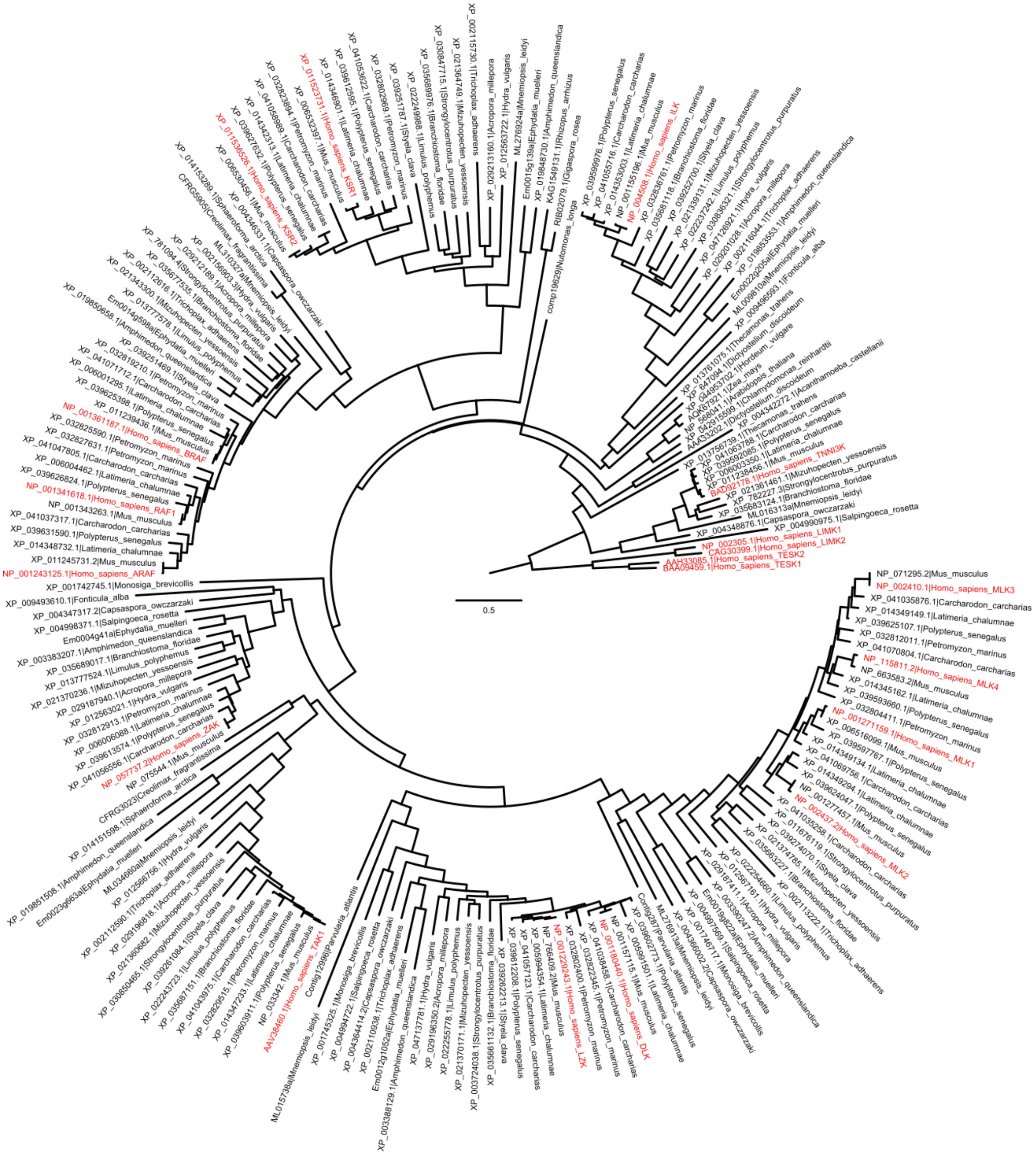


**Fig. S8. Maximum-likelihood TKL MAP3K phylogeny.** The TKL MAP3K phylogeny with the highest likelihood value among the ten technical replicates. The tree is rooted with human outgroup kinases. All human homologs are colored red and labeled with the names of corresponding proteins. Branch length represents the expected number of substitutions per unit time.

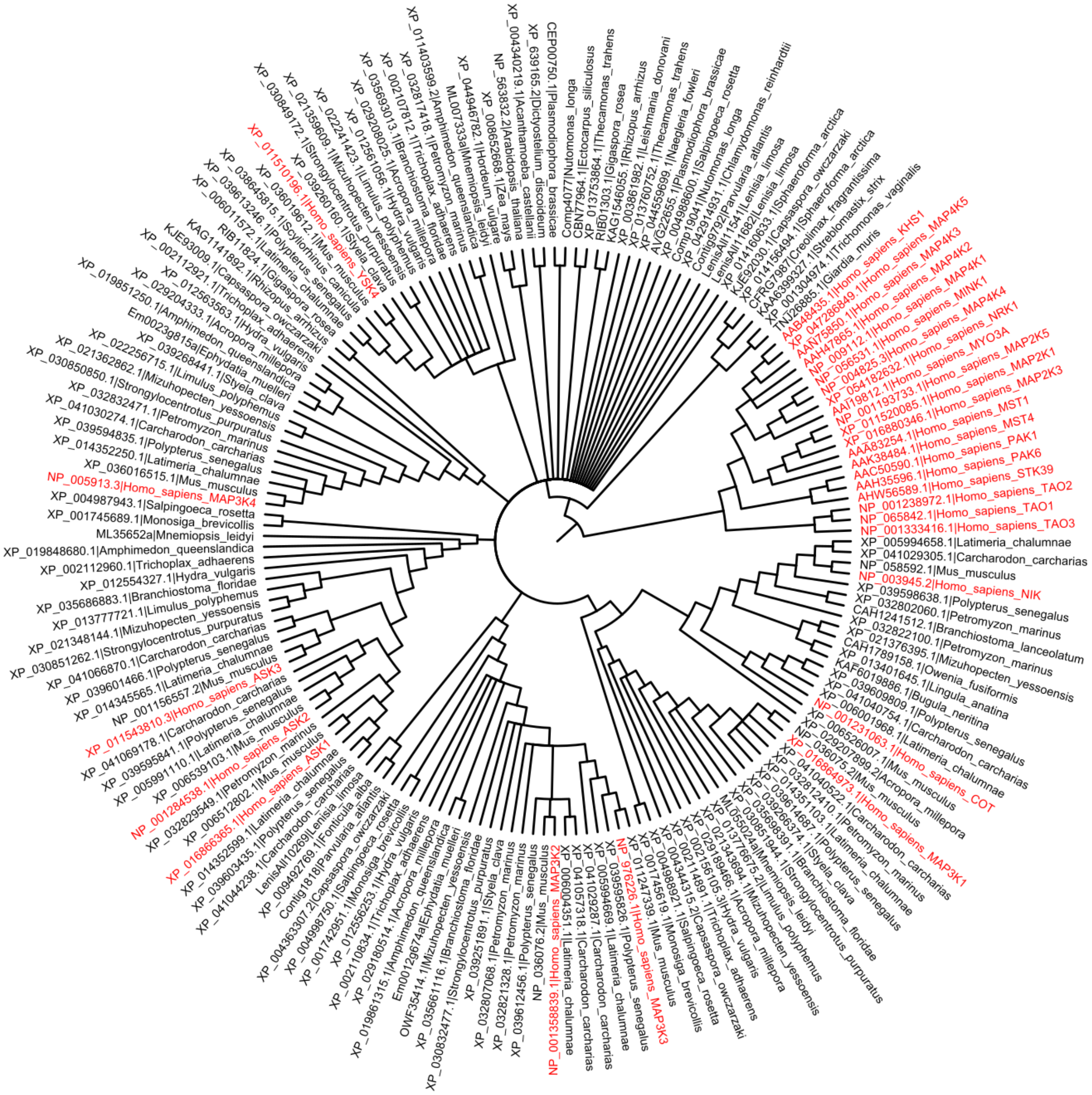

**Fig. S9. Strict consensus STE MAP3K phylogeny.** Strict consensus STE MAP3K cladogram of the ten maximum likelihood reconstruction replicates. The tree is rooted with human outgroup kinases. All human homologs are colored red and labeled with the names of corresponding proteins.

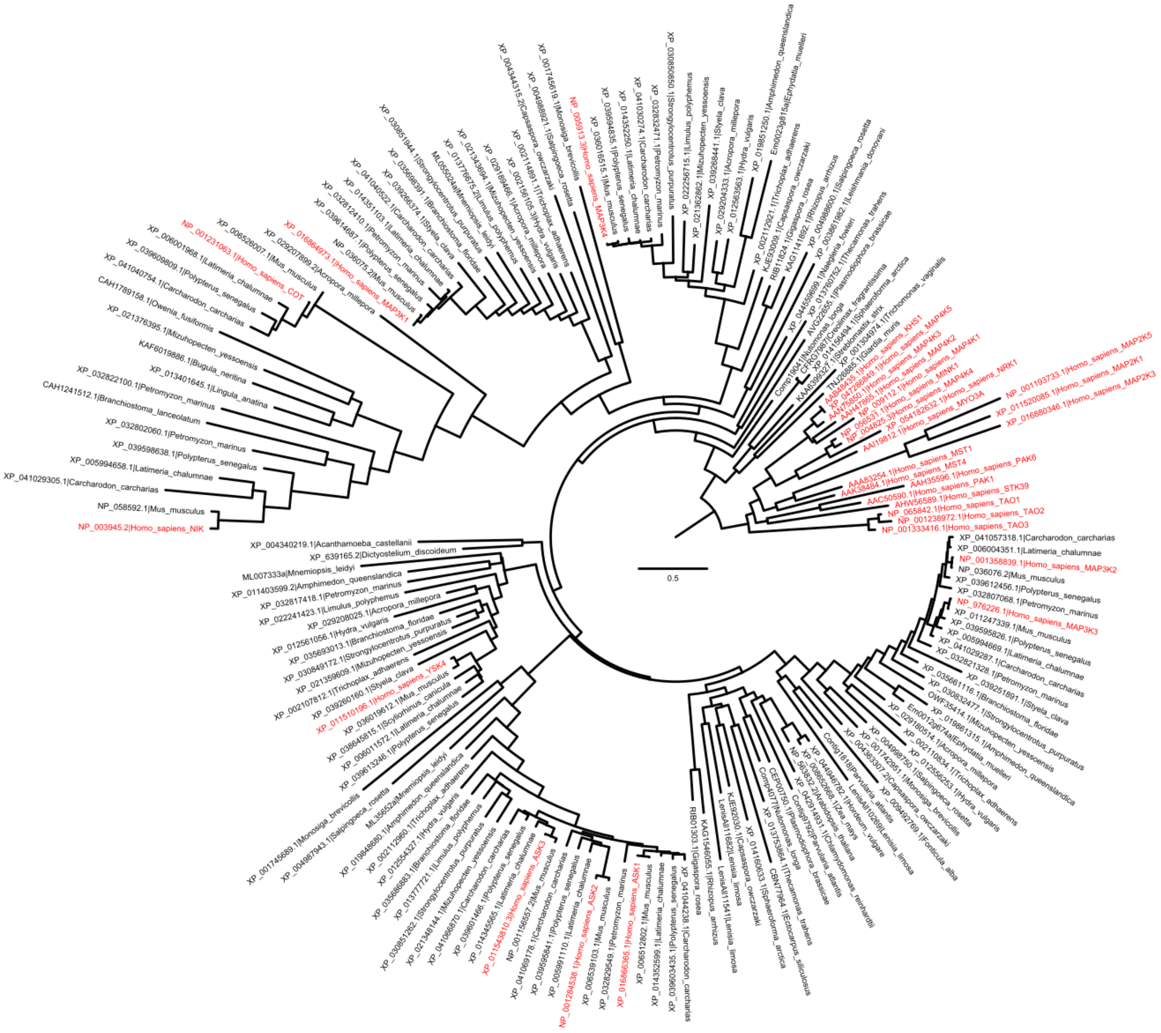


**Fig. S10. Maximum-likelihood STE MAP3K phylogeny.** The STE MAP3K phylogeny with the highest likelihood value among the ten technical replicates. The tree is rooted with human outgroup kinases. All human homologs are colored red and labeled with the names of corresponding proteins. Branch length represents the expected number of substitutions per unit time.

**Table S1. MAPK homologs.** MAPK homologs included in this study. Homologs are cataloged by inferred lineages; those that cannot be confidently identified are listed under unknown. Homologs listed under 'other' are either basal lineages or recovered at position challenging to classified. Homologs listed under 'removed' are removed from this study due to improbable reconstructed position. Taxa with sequences acquired outside from Genbank (Clark et al. 2016) include Ephydatia muelleri (Kenny et al. 2020), Mnemiopsis leidyi (Moreland et al. 2020), Creolimax fragrantissima (de Mendoza et al. 2015), Sphaeroforma arctica (Dudin et al. 2019), Parvularia atlantis (Multicellgenome Lab, 2017a), and Lenisia limosa (Multicellgenome lab, 2017b).

| | MAPK | | | | | | | | | | | | | | | | |
|---|---|---|---|---|---|---|---|---|---|---|---|---|---|---|---|---|---|
| Taxon | ERK1/2 or *MRCA* ERK1/3 | | ERK3/4 | | ERK5 | JNK | | | p38 or *MRCA* JNK/p38 | | | | NLK | ERK7 | other (basal/unlcear) | | removed |
| Homo sapiens | NP_002736.3 | AHW56518.1 | NP_002738.2 | NP_002739.1 | AAH30134.1 | AAI30571.1 | AAH32539.1 | P53779.2 | NP_002742.3 | AAH15741.1 | NP_002745.1 | AAH31574.1 | NP_057315.3 | NP_620590.2 | | | |
| Mus musculus | NP_001033752.1 | NP_036082.1 | NP_766220.2 | NP_056621.4 | NP_001277963.1 | NP_001297382.1 | NP_997575.2 | NP_001305031.1 | NP_035291.4 | NP_038899.1 | NP_036080.2 | NP_036081.1 | NP_032728.3 | NP_808590.1 | | | |
| Latimeria chalumnae | XP_005990239.1 | XP_006005975.1 | XP_006007906.1 | XP_006001969.1 | XP_014349945.1 | XP_014351336.1 | XP_006003530.1 | XP_014346489.1 | XP_014342863.1 | XP_005994304.1 | XP_005995531.2 | XP_006013641.1 | XP_005987613.2 | XP_014346981.1 | | | |
| Polypterus senegalus | XP_039627493.1 | XP_039620893.1 | XP_039606332.1 | XP_039626247.1 | XP_039598485.1 | XP_039630024.1 | XP_039606744.1 | | XP_039625739.1 | XP_039625744.1 | XP_039605224.1 | XP_039605223.1 | XP_039612584.1 | XP_039593111.1 | | | |
| Carcharodon carcharias | XP_041058004.1 | | XP_041049628.1 | XP_041030967.1 | XP_041038075.1 | XP_041032082.1 | XP_041042736.1 | | XP_041059494.1 | XP_041059495.1 | XP_041050843.1 | | XP_041052761.1 | XP_041041090.1 | | | |
| Petromyzon marinus | XP_032803009.1 | | XP_032809784.1 | | XP_032800128.1 | XP_032805849.1 | XP_032825168.1 | | XP_032825624.1 | XP_032810022.1 | XP_032810021.1 | | XP_032823522.1 | XP_032817309.1 | | | |
| Styela clava | XP_039266663.1 | | XP_039253102.1 | | | XP_039253581.1 | | | XP_039264856.1 | | | | XP_039271258.1 | XP_039265837.1 | | | |
| Branchiostoma floridae | XP_035691013.1 | | | | XP_035692155.1 | XP_035669587.1 | | | XP_035676942.1 | | | | XP_035690656.1 | XP_035664664.1 | | | |
| Strongylocentrotus purpuratus | NP_999813.1 | | XP_011669368.2 | | XP_030847564.1 | XP_030835758.1 | | | XP_030827864.1 | | | | XP_003729479.2 | XP_030854361.1 | | | |
| Mizuhopecten yessoensis | ANG60946.1 | | XP_021358514.1 | | XP_021346677.1 | XP_021346752.1 | | | XP_021356955.1 | | | | XP_021374726.1 | XP_021379016.1 | | | |
| Limulus polyphemus | XP_013783258.1 | | | | | XP_022245910.1 | | | XP_013782451.1 | | | | XP_013775523.1 | XP_013792929.2 | | | |
| Acropora millepora | XP_029188819.1 | | XP_029193217.2 | | XP_029194842.1 | XP_029206583.1 | | | XP_029201498.1 | | | | XP_029190737.1 | XP_029212501.1 | | | |
| Hydra vulgaris | XP_002154499.3 | | XP_002158990.1 | | XP_012556163.1 | NP_001352455.1 | | | XP_002156118.2 | | | | XP_004210667.2 | XP_012560480.1 | | | |
| Trichoplax adhaerens | XP_002115346.1 | | | | XP_002108552.1 | XP_002115959.1 | | | XP_002111921.1 | | | | XP_002114970.1 | XP_002114430.1 | | | |
| Amphimedon queenslandica | XP_003388915.1 | | XP_003384372.1 | | XP_003389469.1 | XP_003384215.1 | | | XP_003384011.1 | | | | XP_003387118.2 | XP_003387542.1 | | | |
| Ephydatia muelleri | Em0087g15a | | | | Em0015g921a | Em0002g1177a | | | Em0001g2907a | | | | Em0015g1255a | Em0023g39a | | | |
| Mnemiopsis leidyi | ML00124a | | ML06751a | | | ML08261a | | | ML09425a | | | | | ML205610a | | | |
| Monosiga brevicollis | XP_001748004.1 | | XP_001742351.1 | | XP_001742318.1 | XP_001743472.1 | | | XP_001750951.1 | | | | | | | | |
| Salpingoeca rosetta | | | XP_004988001.1 | | XP_004991146.1 | XP_004988833.1 | | | | | | | | | | | XP_004996292.1 |
| Capsaspora owczarzaki | XP_004342477.2 | | | | XP_004345440.1 | | | | XP_004347920.1 | | | | | | | | XP_004349328.2 |
| Creolimax fragrantissima | CFRG3899 | | | | | | | | CFRG0929 | | | | | | | | CFRG7629 |
| Sphaeroforma arctica | XP_014153190.1 | | | | | | | | XP_014153626.1 | | | | | | | | XP_014150243.1 |
| Rhizopus oryzae | KAG1536847.1 | | | | KAG1140200.1 | | | | KAG1162933.1 | | | | | | | | |
| Gigaspora rosea | RIB17943.1 | | | | RIB18483.1 | | | | RIB04295.1 | | | | | | | | |
| Fonticula alba | | | | | XP_009495058.1 | XP_009494093.1 | | | | | | | | | | | XP_009494434.1 |
| Parvularia atlantis | | | | | Contig3395 | | | | Contig4180 | | | | | | | | |
| Thecamonas trahens | | | | | | | | | | | | | | XP_013756540.1 | XP_013754506.1 | XP_013760386.1 | |
| Lenisia limosa | | | | | | | | | | | | | | LenisAll4658 | LenisAll11816 | LenisAll1389 | |
| Acanthamoeba castellanii | | | | | | | | | | | | | | XP_004333975.1 | XP_004344888.1 | | |
| Dictyostelium discoideum | | | | | | | | | | | | | | XP_638833.1 | AAA59387.1 | | |
| Nutomonas longa | | | | | | | | | | | | | | comp20771 | comp23530 | | |
| Streblomastix strix | | | | | | | | | | | | | | KAA6378207.1 | KAA6403363.1 | | |
| Giardia muris | | | | | | | | | | | | | | TNJ29519.1 | TNJ28019.1 | | |
| Trichomonas vaginalis | | | | | | | | | | | | | | XP_001323663.1 | XP_001323034.1 | | |
| Chlamydomonas reinhardtii | | | | | | | | | | | | | | XP_001699095.2 | XP_001700291.1 | | |
| Hordeum vulgare | | | | | | | | | | | | | | BAK02898.1 | BAJ86277.1 | | |
| Zea mays | | | | | | | | | | | | | | AAS64372.2 | NP_001104843.2 | | |
| Arabidopsis thaliana | | | | | | | | | | | | | | | NP_192046.1 | | |
| Ectocarpus siliculosus | | | | | | | | | | | | | | CBJ27517.1 | CBN78224.1 | | |
| Plasmodiophora brassicae | | | | | | | | | | | | | | AVG22642.1 | AVG22639.1 | | |
| Leishmania donovani | | | | | | | | | | | | | | XP_003865194.1 | XP_003858890.1 | | |
| Naegleria fowleri | | | | | | | | | | | | | | KAF0972656.1 | KAF0982511.1 | | |

**Table S2. MAP2K homologs.** MAP2K homologs included in this study. Homologs are cataloged by inferred lineages; those that cannot be confidently identified are listed under unknown. Homologs listed under 'other' are either basal lineages or recovered at position challenging to classified. Homologs listed under 'removed' are removed from this study due to improbable reconstructed position. Taxa with sequences acquired outside from Genbank (Clark et al. 2016) include Ephydatia muelleri (Kenny et al. 2020), Mnemiopsis leidyi (Moreland et al. 2020), Creolimax fragrantissima (de Mendoza et al. 2015), Sphaeroforma arctica (Dudin et al. 2019), Parvularia atlantis (Multicellgenome Lab, 2017a), and Lenisia limosa (Multicellgenome lab, 2017b).

| | MAP2K | | | | | | | | |
|---|---|---|---|---|---|---|---|---|---|
| Taxon | map2k1/2 | | map2k3/6 or *MRCA* MAPK3/4/7 | | map2k4 | map2k5 | map2k7 | other (basal/unlcear) | | removed |
| Homo sapiens | XP_011520085.1 | XP_016882478.1 | XP_016880346.1 | XP_011523328.1 | NP_001268364.1 | NP_001193733.1 | NP_001284484.1 | | | |
| Mus musculus | XP_006511259.1 | NP_001334073.1 | NP_032954.1 | NP_036073.1 | NP_001303296.1 | NP_035970.1 | NP_001036022.1 | | | |
| Latimeria chalumnae | XP_005990160.1 | XP_005999133.1 | XP_006008481.1 | XP_005992212.2 | XP_006011478.1 | XP_006001847.1 | XP_014347656.1 | | | |
| Polypterus senegalus | XP_039629458.1 | XP_039622866.1 | XP_039632221.1 | | XP_039595092.1 | XP_039629441.1 | XP_039626376.1 | | | |
| Carcharodon carcharias | XP_041031084.1 | XP_041060212.1 | XP_041062215.1 | XP_041072910.1 | XP_041072798.1 | XP_041030662.1 | XP_041033301.1 | | | |
| Petromyzon marinus | XP_032829743.1 | XP_032804312.1 | XP_032825980.1 | XP_032830445.1 | XP_032816932.1 | XP_032804405.1 | XP_032802591.1 | | | |
| Styela clava | XP_039257658.1 | | XP_039263764.1 | | XP_039263192.1 | XP_039254583.1 | XP_039269677.1 | | | |
| Branchiostoma floridae | XP_035665490.1 | | XP_035695198.1 | | XP_035696000.1 | XP_035666863.1 | XP_035683536.1 | | | |
| Strongylocentrotus purpuratus | XP_011677450.1 | | XP_030840761.1 | | XP_030840399.1 | XP_030832379.1 | XP_030853441.1 | | | |
| Mizuhopecten yessoensis | XP_021372354.1 | | XP_021376386.1 | | XP_021341288.1 | XP_021362175.1 | AKS30508.1 | | | |
| Limulus polyphemus | XP_022253276.1 | | XP_013784770.1 | | XP_013781038.1 | | XP_013786818.2 | | | |
| Acropora millepora | XP_029192343.1 | | XP_029193377.1 | | XP_029192273.1 | XP_029200799.1 | XP_029180030.1 | | | |
| Hydra vulgaris | XP_002161947.2 | | XP_012562177.1 | | XP_002162501.3 | XP_012558557.1 | XP_002162140.3 | | | |
| Trichoplax adhaerens | RDD46366.1 | | RDD43980.1 | | XP_002109210.1 | RDD40670.1 | RDD43374.1 | | | |
| Amphimedon queenslandica | XP_003384864.1 | | XP_003384504.1 | | XP_011410455.1 | | XP_019851593.1 | | | |
| Ephydatia muelleri | Em0009g969a | | Em0020g172a | | Em0020g81a | | | | | |
| Mnemiopsis leidyi | ML02994a | | ML070266a | | ML017945a | | ML07766a | | | |
| Monosiga brevicollis | XP_001745338.1 | | XP_001747273.1 | | | XP_001745245.1 | | | | |
| Salpingoeca rosetta | XP_004994653.1 | | XP_004989160.1 | | | XP_004994592.1 | | | | |
| Capsaspora owczarzaki | KJE90307.1 | | XP_004346163.1 | | | XP_004365166.2 | | | | KJE88420.1 |
| Creolimax fragrantissima | CFRG7646 | | CFRG6538 | | | | | | | CFRG5305 |
| Sphaeroforma arctica | Sarc4_g30182T | | XP_014158900.1 | | | | | XP_014156621.1 | | XP_014156622.1 |
| Rhizopus oryzae | KAG1542273.1 | | KAG1211502.1 | | | KAG0743027.1 | | | | |
| Gigaspora rosea | RIB14944.1 | | RIB11085.1 | | | RIB10023.1 | | | | |
| Fonticula alba | XP_009494286.1 | | | | | | | | | XP_009492389.1 |
| Parvularia atlantis | Contig7293 | | Contig3416 | | | | | | | Contig6469 |
| Thecamonas trahens | XP_013761493.1 | | XP_013760746.1 | | | XP_013758939.1 | | | | |
| Lenisia limosa | | | | | | | | LenisAll8332 | | |
| Acanthamoeba castellanii | | | | | | | | XP_004333955.1 | | |
| Dictyostelium discoideum | | | | | | | | AAB58577.1 | | |
| Nutomonas longa | | | | | | | | comp12234 | comp23822 | |
| Streblomastix strix | | | | | | | | KAA6397582.1 | | |
| Giardia muris | | | | | | | | TNJ26678.1 | | |
| Trichomonas vaginalis | | | | | | | | XP_001318307.1 | | |
| Chlamydomonas reinhardtii | | | | | | | | XP_001696437.1 | | |
| Hordeum vulgare | | | | | | | | BAU62327.1 | | |
| Zea mays | | | | | | | | XP_008672206.1 | | |
| Arabidopsis thaliana | | | | | | | | NP_001318713.1 | | |
| Ectocarpus siliculosus | | | | | | | | CBN78311.1 | | |
| Plasmodiophora brassicae | | | | | | | | AVG22647.1 | | |
| Leishmania donovani | | | | | | | | AYU83436.1 | | |
| Naegleria fowleri | | | | | | | | XP_044564510.1 | | |

**Table S3. TKL MAP3K homologs.** TKL MAP3K homologs included in this study. Homologs are cataloged by inferred lineages; those that cannot be confidently identified are listed under unknown. Homologs listed under 'other' are either basal lineages or recovered at position challenging to classified. Taxa with sequences acquired outside from Genbank (Clark et al. 2016) include Ephydatia muelleri (Kenny et al. 2020), Mnemiopsis leidyi (Moreland et al. 2020), Creolimax fragrantissima (de Mendoza et al. 2015), Sphaeroforma arctica (Dudin et al. 2019), Parvularia atlantis (Multicellgenome Lab, 2017a), and Lenisia limosa (Multicellgenome lab, 2017b).

| | TKL | | | | | | | | | | | | | | | |
|---|---|---|---|---|---|---|---|---|---|---|---|---|---|---|---|---|
| Taxon | MLK1/2/3/4 | | | | DLK/LZK | | TAK1 | ZAK | ILK | TNNI3K | KSR1/2 | | ARAF/BRAF/RAF1 | | | other (basal/unlcear) |
| Homo sapiens | NP_001271159.1 | NP_002437.2 | NP_002410.1 | NP_115811.2 | NP_001180440.1 | NP_001229243.1 | AAV38460.1 | NP_057737.2 | NP_004508.1 | BAD92178.1 | XP_011523731.1 | XP_011536526.1 | NP_001243125.1 | NP_001361187.1 | NP_001341618.1 | |
| Mus musculus | XP_006516099.1 | NP_001277457.1 | NP_071295.2 | NP_663583.2 | NP_001157115.1 | NP_766409.2 | NP_033342.1 | NP_075544.1 | NP_001155196.1 | XP_011238456.1 | XP_006532397.1 | XP_006530456.1 | XP_011245731.2 | XP_011239436.1 | NP_001343263.1 | |
| Latimeria chalumnae | XP_014349134.1 | XP_014349294.1 | XP_014349149.1 | XP_014345162.1 | XP_005991501.1 | XP_005994354.1 | XP_014347233.1 | XP_006006088.1 | XP_014353003.1 | XP_006003350.1 | XP_014346901.1 | XP_014342313.1 | XP_014348732.1 | XP_006001295.1 | XP_006004462.1 | |
| Polypterus senegalus | XP_039597767.1 | XP_039624047.1 | XP_039625107.1 | XP_039593660.1 | XP_039602773.1 | XP_039612208.1 | XP_039603911.1 | XP_039613574.1 | XP_039599976.1 | XP_039592085.1 | XP_039612595.1 | XP_039627632.1 | XP_039631590.1 | XP_039625398.1 | XP_039626824.1 | |
| Carcharodon carcharias | XP_041069756.1 | XP_041035258.1 | XP_041035876.1 | XP_041070804.1 | XP_041038458.1 | XP_041057123.1 | XP_041043975.1 | XP_041056556.1 | XP_041055716.1 | XP_041063788.1 | XP_041053622.1 | XP_041058989.1 | XP_041037317.1 | XP_041071712.1 | XP_041047805.1 | |
| Petromyzon marinus | XP_032812011.1 | XP_032804411.1 | | | XP_032802400.1 | XP_032822345.1 | XP_032829515.1 | XP_032812913.1 | XP_032836761.1 | | XP_032802969.1 | XP_032823894.1 | XP_032827631.1 | XP_032819210.1 | XP_032825590.1 | |
| Styela clava | XP_039274070.1 | | | | XP_039262213.1 | | XP_039261084.1 | | XP_039252700.1 | | XP_039251787.1 | | XP_039251469.1 | | | |
| Branchiostoma floridae | XP_035663227.1 | | | | XP_035661132.1 | | XP_035687151.1 | XP_035689017.1 | XP_035681118.1 | XP_035683124.1 | XP_035689976.1 | | XP_035677535.1 | | | |
| Strongylocentrotus purpuratus | XP_011676119.1 | | | | XP_003724038.1 | | XP_030850465.1 | | XP_030836321.1 | XP_782227.3 | XP_030847715.1 | | XP_781094.4 | | | |
| Mizuhopecten yessoensis | XP_021374785.1 | | | | XP_021370171.1 | | XP_021360682.1 | XP_021370236.1 | XP_021339131.1 | XP_021361461.1 | XP_021364749.1 | | XP_021343300.1 | | | |
| Limulus polyphemus | XP_022254660.1 | | | | XP_022255778.1 | | XP_022243723.1 | XP_013777524.1 | XP_022237242.1 | | XP_022249988.1 | | XP_013777578.1 | | | |
| Acropora millepora | XP_029187411.1 | | | | XP_029196350.2 | | XP_029196818.1 | XP_029187940.1 | XP_029201028.1 | | XP_029213160.1 | | XP_029212189.1 | | | |
| Hydra vulgaris | XP_012567161.1 | | | | XP_047137781.1 | | XP_012566756.1 | XP_012563021.1 | XP_047126921.1 | | XP_012563722.1 | | XP_002156903.3 | | | |
| Trichoplax adhaerens | XP_002113222.1 | | | | XP_002110938.1 | | XP_002112590.1 | | XP_002116044.1 | | XP_002115730.1 | | XP_002112616.1 | | | |
| Amphimedon queenslandica | XP_003390247.2 | | | | XP_003388129.1 | | XP_019851508.1 | XP_003383207.1 | XP_019853553.1 | | XP_019848730.1 | | XP_019850658.1 | | | |
| Ephydatia muelleri | Em0019g822a | | | | Em0012g1052a | | Em0023g663a | Em0004g41a | Em0022g205a | | Em0015g139a | | Em0014g598a | | | |
| Mnemiopsis leidyi | ML276913a | | | | ML015738a | | ML034660a | | ML009810a | ML016313a | ML276924a | | ML310327a | | | |
| Monosiga brevicollis | XP_001746717.1 | | | | XP_001745325.1 | | | | | | | | | | | XP_001742745.1 |
| Salpingoeca rosetta | XP_004997569.1 | | | | XP_004994722.1 | | | XP_004998371.1 | | XP_004990975.1 | | | | | | |
| Capsaspora owczarzaki | XP_004366002.2 | | | | XP_004364414.2 | | | XP_004347317.2 | | XP_004348876.1 | | | XP_004346331.1 | | | |
| Creolimax fragrantissima | | | | | | | CFRG3023 | | | | | | CFRG5905 | | | |
| Sphaeroforma arctica | | | | | | | XP_014151598.1 | | | | | | XP_014153289.1 | | | |
| Rhizopus oryzae | | | | | | | | | | | | | | | | KAG1549131.1 |
| Gigaspora rosea | | | | | | | | | | | | | | | | RIB02079.1 |
| Fonticula alba | | | | | | | | XP_009493610.1 | XP_009496593.1 | | | | | | | |
| Parvularia atlantis | Contig287 | | | | Contig12996 | | | | | | | | | | | |
| Thecamonas trahens | | | | | | | | | XP_013761075.1 | | | | | | | XP_013756739.1 |
| Lenisia limosa | | | | | | | | | | | | | | | | |
| Acanthamoeba castellanii | | | | | | | | | | | | | | | | XP_004342272.1 |
| Dictyostelium discoideum | | | | | | | | | XP_647094.1 | | | | | | | AAA33202.1 |
| Nutomonas longa | | | | | | | | | | | | | | | | comp19629 |
| Streblomastix strix | | | | | | | | | | | | | | | | |
| Giardia muris | | | | | | | | | | | | | | | | |
| Trichomonas vaginalis | | | | | | | | | | | | | | | | |
| Chlamydomonas reinhardtii | | | | | | | | | | | | | | | | XP_042915599.1 |
| Hordeum vulgare | | | | | | | | | | | | | | | | XP_044953702.1 |
| Zea mays | | | | | | | | | | | | | | | | AQK67921.1 |
| Arabidopsis thaliana | | | | | | | | | | | | | | | | NP_568041.1 |
| Ectocarpus siliculosus | | | | | | | | | | | | | | | | |
| Plasmodiophora brassicae | | | | | | | | | | | | | | | | |
| Leishmania donovani | | | | | | | | | | | | | | | | |
| Naegleria fowleri | | | | | | | | | | | | | | | | |

**Table S4. STE MAP3K homologs.** STE MAP3K homologs included in this study. Homologs are cataloged by inferred lineages; those that cannot be confidently identified are listed under unknown. Homologs listed under 'other' are either basal lineages or recovered at position challenging to classified. Homolog listed under 'additional' are orthologs of MRCACOT/NIK and included for enhancing representation of the COT/NIK lineage. Taxa with sequences acquired outside from Genbank (Clark et al. 2016) include Ephydatia muelleri (Kenny et al. 2020), Mnemiopsis leidyi (Moreland et al. 2020), Creolimax fragrantissima (de Mendoza et al. 2015), Sphaeroforma arctica (Dudin et al. 2019), Parvularia atlantis (Multicellgenome Lab, 2017a), and Lenisia limosa (Multicellgenome lab, 2017b).

| | STE | | | | | | | | | | | |
|---|---|---|---|---|---|---|---|---|---|---|---|---|
| Taxon | MAP3K1 or *MRCA* MAP3K1/COT | COT/NIK | | MAP3K2/3 | | MAP3K4 | ASK1/2/3 | | | YSK4 | other (basal/unlcear) | |
| Homo sapiens | XP_016864973.1 | NP_001231063.1 | NP_003945.2 | NP_001358839.1 | NP_976226.1 | NP_005913.3 | XP_016866365.1 | NP_001284538.1 | XP_011543810.3 | XP_011510196.1 | | |
| Mus musculus | NP_036075.2 | XP_006526007.1 | NP_058592.1 | NP_036076.2 | XP_011247339.1 | XP_036016515.1 | XP_006512802.1 | XP_006539103.1 | NP_001156557.2 | XP_036019612.1 | | |
| Latimeria chalumnae | XP_014351103.1 | XP_006001968.1 | XP_005994658.1 | XP_006004351.1 | XP_005994669.1 | XP_014352250.1 | XP_014352599.1 | XP_005991110.1 | XP_014345565.1 | XP_006011572.1 | | |
| Polypterus senegalus | XP_039614687.1 | XP_039609809.1 | XP_039598638.1 | XP_039612456.1 | XP_039595826.1 | XP_039594835.1 | XP_039603435.1 | XP_039595841.1 | XP_039601466.1 | XP_039613246.1 | | |
| Carcharodon carcharias | XP_041040522.1 | XP_041040754.1 | XP_041029305.1 | XP_041057318.1 | XP_041029287.1 | XP_041030274.1 | XP_041044238.1 | XP_041069178.1 | XP_041066870.1 | XP_038645815.1 (Scyliorhinus_canicula) | | |
| Petromyzon marinus | XP_032812410.1 | XP_032822100.1 | XP_032802060.1 | XP_032807068.1 | XP_032821328.1 | XP_032832471.1 | XP_032829549.1 | | | XP_032817418.1 | | |
| Styela clava | XP_039266374.1 | | | XP_039251891.1 | | XP_039268441.1 | | | | XP_039260160.1 | | |
| Branchiostoma floridae | XP_035698391.1 | CAH1241512.1 | | XP_035661116.1 | | | XP_035686883.1 | | | XP_035693013.1 | | |
| Strongylocentrotus purpuratus | XP_030851944.1 | | | XP_030832477.1 | | XP_030850850.1 | XP_030851262.1 | | | XP_030849172.1 | | |
| Mizuhopecten yessoensis | XP_021343694.1 | XP_021376395.1 | | OWF35414.1 | | XP_021362862.1 | XP_021348144.1 | | | XP_021359609.1 | | |
| Limulus polyphemus | XP_013776675.2 | | | | | XP_022256715.1 | XP_013777721.1 | | | XP_022241423.1 | | |
| Acropora millepora | XP_029189466.1 | XP_029207899.2 | | XP_029180514.1 | | XP_029204333.1 | | | | XP_029208025.1 | | |
| Hydra vulgaris | XP_002156105.3 | | | XP_012556253.1 | | XP_012563563.1 | XP_012554327.1 | | | XP_012561056.1 | | |
| Trichoplax adhaerens | XP_002114891.1 | additional | | XP_002110834.1 | | XP_002112921.1 | XP_002112960.1 | | | XP_002107812.1 | | |
| Amphimedon queenslandica | | Bugula_neritina | KAF6019886.1 | XP_019861315.1 | | XP_019851250.1 | XP_019848680.1 | | | XP_011403599.2 | | |
| Ephydatia muelleri | | Owenia_fusiformis | CAH1789158.1 | Em0012g674a | | Em0023g815a | | | | | | |
| Mnemiopsis leidyi | ML055024a | Lingula_anatina | XP_013401645.1 | ML038027a | | | ML35652a | | | ML007333a | | |
| Monosiga brevicollis | XP_001745619.1 | | | XP_001742951.1 | | | XP_001745689.1 | | | | | |
| Salpingoeca rosetta | XP_004988921.1 | | | XP_004998750.1 | | | XP_004987943.1 | | | | | |
| Capsaspora owczarzaki | XP_004344315.2 | | | XP_004363307.2 | | KJE93009.1 | | | | | KJE92030.1 | |
| Creolimax fragrantissima | | | | | | | | | | | CFRG7987 | |
| Sphaeroforma arctica | | | | | | | | | | | XP_014160633.1 | XP_014156494.1 |
| Rhizopus oryzae | | | | | | KAG1141892.1 | | | | | KAG1546055.1 | |
| Gigaspora rosea | | | | | | RIB11824.1 | | | | | RIB01303.1 | |
| Fonticula alba | | | | XP_009492769.1 | | | | | | | | |
| Parvularia atlantis | | | | Contig1818 | | | | | | | Contig9792 | |
| Thecamonas trahens | | | | | | | | | | | XP_013760752.1 | XP_046160633.1 |
| Lenisia limosa | | | | LenisAll10269 | | | | | | | LenisALL11682 | LenisAll11541 |
| Acanthamoeba castellanii | | | | | | | | | | | XP_004340219.1 | |
| Dictyostelium discoideum | | | | | | | | | | | XP_639165.2 | |
| Nutomonas longa | | | | | | | | | | | Comp19041 | Comp4077 |
| Streblomastix strix | | | | | | | | | | | KAA6399327.1 | |
| Giardia muris | | | | | | | | | | | TNJ26885.1 | |
| Trichomonas vaginalis | | | | | | | | | | | | |
| Chlamydomonas reinhardtii | | | | | | | | | | | XP_042914931.1 | |
| Hordeum vulgare | | | | | | | | | | | XP_044946782.1 | |
| Zea mays | | | | | | | | | | | XP_008652668.1 | |
| Arabidopsis thaliana | | | | | | | | | | | NP563832.2 | |
| Ectocarpus siliculosus | | | | | | | | | | | CBN77964.1 | |
| Plasmodiophora brassicae | | | | | | | | | | | AVG22655.1 | CEP00750.1 |
| Leishmania donovani | | | | | | | | | | | XP_003861921.1 | |
| Naegleria fowleri | | | | | | | | | | | XP_044559699.1 | |

### SI References


1. Clark, K., Karsch-Mizrachi, I., Lipman, D. J., Ostell, J., & Sayers, E. W. (2016). GenBank. Nucleic Acids Research, 44, D67-D72. https://doi.org/10.1093/nar/gkv1276
2. Kenny, N. J., Francis, W. R., Rivera-Vicéns, R. E., Juravel, K., de Mendoza, A., Díez-Vives, C., Lister, R., Bezares-Calderón, L. A., Grombacher, L., Roller, M., Barlow, L. D., Camilli, S., Ryan, J. F., Wörheide, G., Hill, A. L., Riesgo, A., & Leys, S. P. (2020). Tracing animal genomic evolution with the chromosomal-level assembly of the freshwater sponge Ephydatia muelleri. Nature Communications, 11(1), 3676. https://doi.org/10.1038/s41467-020-17397-w
3. Moreland, R. T., Nguyen, A., Ryan, J. F., & Baxevanis, A. D. (2020). The Mnemiopsis genome project portal: Integrating new gene expression resources and improving data visualization. Database: The Journal of Biological Databases and Curation, 2020. https://doi.org/10.1093/database/baaa029
4. de Mendoza, A., Suga, H., Permanyer, J., Irimia, M., & Ruiz-Trillo, I. (2015). Complex transcriptional regulation and independent evolution of fungal-like traits in a relative of animals. eLife, 4, e08904. https://doi.org/10.7554/eLife.08904
5. Dudin, O., Ondracka, A., Grau-Bové, X., Haraldsen, A. A., Toyoda, A., Suga, H., Bråte, J., & Ruiz-Trillo, I. Transcriptome - Sphaeroforma arctica. https://doi.org/10.6084/m9.figshare.8299529.v2
6. Multicellgenome Lab, & Torruella, G. (2017a). Transcriptome - Parvularia atlantis. https://doi.org/10.6084/m9.figshare.3898485.v4
7. Multicellgenome Lab, & Torruella, G. (2017b). Transcriptome - *Nutomonas longa*. https://doi.org/10.6084/m9.figshare.4560862.v2
